# The circ_0074371/miR-4738-3p/AMPK Signaling Axis Induces M1 Macrophage Polarization and Restricts Intracellular Survival of Mycobacterium tuberculosis

**DOI:** 10.64898/2026.10.20.760595

**Authors:** Jun-Ai Zhang, Chen-Ye Lian, Yuan-Bin Lu, Shu-Ying Yao, Huan Xu, Yao Xu, Gan-Bing Liu, Guo-Si Chen, Yue-Feng Cen, Yuan-Ting Li, Zhuo-Lin Xu, Yi-Dan Huang, Guo-Ying Chen, Jia-Jun Wang, Jia-Hua Shi, Yu-Chi Gao, Xiao-Zhen Cai, Jing Liang, Wen-Cai Huang, Zhi-Ping Zhang, Jiang Pi, Jun-Fa Xu

## Abstract

Tuberculosis, an infectious disease caused by *Mycobacterium tuberculosis*, remains endemic in regions where it is prevalent. *Mycobacterium tuberculosis* is an intracellular pathogen that can survive within host macrophages; however, the mechanisms governing its long-term persistence and the progression to active pulmonary tuberculosis remain poorly understood. Circular RNAs (circRNAs) have been implicated in the initiation and progression of numerous diseases. In our previous study, circRNA sequencing identified several circRNAs associated with tuberculosis progression. In the present study, sequencing and reverse transcription–polymerase chain reaction (RT-PCR) analyses revealed that Circ_0074371 was significantly downregulated in monocytes from patients with active pulmonary tuberculosis (ATB) and was closely associated with disease onset and progression, exhibiting considerable diagnostic value for ATB. Functional experiments further showed that Circ_0074371 significantly impaired the intracellular survival of *Mtb* and promoted M1 macrophage polarization. Mechanistically, Circ_0074371 acted as a competing endogenous RNA by sequestering miR-4738-3p, a potent negative regulator of AMP-activated protein kinase (AMPK) activation. Collectively, these findings identify Circ_0074371 as a tuberculosis-associated circRNA that regulates AMPK activity and reveal a novel mechanism through which Circ_0074371 influences macrophage polarization via the AMPK pathway, providing new insights into potential therapeutic targets for ATB.

## 1 Introduction

Tuberculosis (TB) is a highly infectious disease caused by *Mycobacterium tuberculosis(MTB)*, which remains the leading cause of death from a single infectious agent, accounting for ∼1.5 million deaths worldwide annually^[1]^. Current TB prevention and control measures remain inadequate, primarily because of the emergence of multidrug-resistant TB^[2]^; he limited efficacy of *Bacillus Calmette–Guérin (BCG)* vaccination in adults, difficulties in diagnosing latent TB infection^[3]^, and HIV coinfection^[4]^. Host-directed immunotherapy has therefore attracted considerable attention as an alternative strategy for TB treatment because its efficacy is not directly affected by bacterial drug resistance^[5]^. During *MTB* infection, macrophages serve as the primary “battlefield” against *MTB*^[6]^,which subsequently activate various immune defense mechanisms of macrophages, such as promoting autophagy, apoptosis and antibacterial M1 polarization, while inhibiting necrosis, ferroptosis, and M2 macrophage polarization, to limit and eradicate the intracellular *MTB* pathogens. However, *MTB* employs a variety of sophisticated strategies to evade host immune clearance and persist within macrophages. These strategies include inhibiting host-cell DNA repair ^[7]^, disrupting lysosomal function^[8]^, and lipid metabolism^[9]^, reprogramming host-cell signaling pathways ^[10]^, and promoting M2 macrophage polarization^[11–14]^. Nevertheless, the underlying molecular mechanisms remain poorly understood, particularly the key regulatory mechanisms governing macrophage polarization.

The role of the AMP-activated protein kinase (AMPK) signaling pathway in macrophage-mediated anti-TB immunity remains insufficiently investigated. Recent studies have shown that activation of AMPK signaling promotes macrophage autophagy and enhances host defense against *MTB*^[15]^,In addition, AMPK cooperates with the mechanistic target of rapamycin (mTOR) pathway to regulate autophagy and cell death during *MTB* infection^[16]^。Although few studies have examined AMPK-mediated regulation of macrophage polarization in TB models, evidence from other disease models indicates that AMPK inhibition promotes M1 macrophage polarization and inflammatory responses^[17]^,This finding is consistent with the observation that AMPK activation suppresses nuclear factor kappa B (NF-κB) signaling^[18]^ 。The AMPK pathway is also involved in macrophage energy sensing^[19–21]^, mitochondrial dysfunction^[22]^, and lipid metabolism^[23]^,Although the direct effects of AMPK deficiency or overexpression on *MTB* infection have not yet been established in vivo or in vitro, AMPK plays a critical regulatory role in macrophage polarization and may represent a key molecular target in host-directed anti-TB therapy.

Nucleic acid therapeutics are rapidly emerging as a transformative therapeutic modality^[24]^,Compared with protein-based molecules, circular RNAs (circRNAs) exhibit greater stability and resistance to degradation, making them promising candidates for the development of anti-TB therapeutics and biomarkers^[25]^,Most annotated circRNAs are generated through the back-splicing of precursor messenger RNAs, in which a downstream 5′ splice site (splice donor) is joined to an upstream 3′ splice site (splice acceptor). This process produces a covalently closed RNA molecule containing a 3′–5′ phosphodiester bond at the back-splice junction (BSJ)^[26]^. CircRNAs regulate diverse cellular processes through multiple mechanisms, including modulation of alternative RNA splicing, cis-regulation of transcription, activity as competitive endogenous RNAs (ceRNAs), translation into peptides, and interaction with microRNAs (miRNAs) to influence their regulatory functions. For example, circRNA ZNF277 acts as a sponge for miR-378d, thereby inhibiting the intracellular survival of *MTB*^[27]^, whereas circTRAPPC6B suppresses *MTB* growth in macrophages and induces autophagy by targeting miR-874-3p^[28]^. Recent studies have demonstrated that circRNAs play important roles in regulating anti-TB immunity^[27,29–32]^,In our previous study, circRNA sequencing identified multiple differentially expressed circRNAs in patients with TB^[25]^, suggesting their potential as TB biomarkers with significant immunoregulatory functions^[33,34]^. Among these circRNAs, circ_0074371 exhibited marked immunomodulatory activity in macrophages and was specifically differentially expressed in monocytes/macrophages. These findings indicate that circ_0074371 has considerable potential for the diagnosis and treatment of TB.

## 2 Materials and methods

### 2.1 Subjects and ethics statement

Peripheral blood samples were collected from patients with active tuberculosis (ATB) at Dongguan Sixth People’s Hospital between March 2019 and September 2020. Patients were eligible for inclusion if they met the diagnostic criteria for tuberculosis outlined in New Tuberculosis by Peng Sanitation, had no history of diabetes, nephritis, hepatitis, HIV infection, or autoimmune disease, and had not recently received immunomodulatory therapy. Healthy controls (HCs) were recruited based on the following criteria: normal chest radiographic findings, no recent history of contact with patients with tuberculosis, a negative tuberculin skin test (TST), and no abnormalities in routine blood test results. The study protocol was approved by the Ethics Committee of Guangdong Medical University, and written informed consent was obtained from all participants. Detailed information on the samples is provided in Appendix III.

### 2.2 Patients and cell samples collection

A total of 48 patients with active pulmonary tuberculosis (ATB), diagnosed as previously described^[35]^, were recruited from Dongguan 6th People’s Hospital, Dongguan, China. Twenty-one healthy volunteers with no bacteriological or clinical evidence of tuberculosis served as healthy controls (HCs) (Table S1, Table S2 and Table S3). All HCs had documented Bacille Calmette–Guérin (*BCG*) vaccination at birth. Participants were enrolled at Dongguan 6th People’s Hospital between December 2019 and September 2020. Individuals with HIV infection, diabetes, cancer, autoimmune disease, a history of immunosuppressive treatment, or previous pulmonary tuberculosis, as described previously^[35]^. were excluded. Sputum samples from all participants were examined by smear microscopy and Ziehl–Neelsen acid-fast staining to determine smear positivity, and were cultured on Lowenstein–Jensen medium according to standard procedures. The individualized treatment regimens for patients with ATB have been described previously^[35]^. Patients initially received anti-tuberculosis drug (ATD) therapy, including the M0 phase (0–4 days) and the M1 phase (25–35 days), with progression to the latter phase defined by the absence of symptoms suggestive of tuberculosis relapse. No significant differences in age or sex were observed between the ATB and HC groups. The study protocol was approved by the Institutional Review and Ethics Boards of Guangdong Medical University and Dongguan 6th People’s Hospital. Written informed consent was obtained from all participants before enrollment.

### 2.3 Cell lines and Mycobacterium

#### 2.3.1 Cell Lines and Primary Cells

The THP-1 cell line is a human peripheral blood monocytic cell line established from the monocytes of a patient with acute monocytic leukemia (ATCC, #TIB-202). Cells were maintained in RPMI 1640 basal medium (Solarbio, China) supplemented with 10% fetal bovine serum (Gibco, USA). To induce differentiation into M0 macrophages in vitro, THP-1 cells were treated with phorbol 12-myristate 13-acetate (PMA; MedChemExpress) at a final concentration of 100 ng/mL for 24 h. The resulting macrophages were primarily used to establish an intracellular *Mycobacterium tuberculosis* (*MTB*) infection model.

To generate a THP-1 cell line overexpressing circ_0074371, the 253-bp sequence corresponding to hsa_circ_0074371 (circBank ID: hsa_circARHGAP26_020) was amplified by PCR. The amplified fragment was digested with EcoRI and BamHI and subsequently ligated into the pLC5-ciR vector. The resulting lentiviral expression plasmid was used to generate lentiviral particles. After appropriate dilution, the lentiviral supernatant was mixed with THP-1 cells and transferred to 24-well plates. The plates were centrifuged at 900–1500 ×g at room temperature for 1 h and then incubated for an additional hour in a CO₂ incubator to promote infection. At 48 h post-infection, the proportion of fluorescent cells was examined by fluorescence microscopy. Cells from the 24-well plates were then distributed into several wells (3–5 wells) and cultured overnight. Puromycin selection was performed at a concentration of 5 µg/mL. Following 24–48 h of selection, cell viability was assessed to determine the optimal puromycin concentration, and cells from the selected wells were used in subsequent experiments. Once the cells reached confluence, they were expanded. GFP-positive cells were isolated by flow cytometry and further expanded. After several passages, the expression of circ_0074371 was verified by quantitative reverse-transcription PCR (qRT-PCR).

For circ_0074371 knockdown, siRNAs targeting the back-splice junction of circ_0074371 were designed and synthesized by Ruibo Biological Technology Co., Ltd. (Guangzhou, China). THP-1 cells were seeded in 12-well plates at a density of 2 × 10⁵ cells per well and cultured overnight. The cells were subsequently treated with PMA for 24 h to induce adherence, reaching approximately 70–80% confluence. For transfection, siRNA was diluted to a final concentration of 50 nM in 30 µL of 1× riboFECT™ CP Buffer, followed by the addition of 3 µL riboFECT™ CP Reagent. The mixture was gently vortexed and incubated at room temperature for 10–15 min. The resulting transfection complexes were then added to the cells. Knockdown efficiency was evaluated by real-time quantitative PCR based on circ_0074371 expression, and the most effective siRNA was selected for subsequent experiments.

The 293T cell line is a human embryonic kidney epithelial cell line derived from human embryonic kidney cells (ATCC). The cells were transfected with the adenovirus E1A gene and express the SV40 large T antigen, including the SV40 origin of replication and promoter region. They were primarily used for dual-luciferase reporter assays and transfection with viral expression vectors.

Peripheral blood mononuclear cells (PBMCs) were isolated as previously described^[36]^. Briefly, approximately 5 mL of peripheral blood was collected from each subject into blood collection tubes containing acid citrate dextrose (ACD). The samples were centrifuged at 1500 rpm for 5 min, after which the plasma was collected and stored at −80°C until analysis. PBMCs were freshly isolated from the blood by standard Ficoll density-gradient centrifugation (GE Healthcare). Cell viability was assessed by trypan blue exclusion and exceeded 95% in all experiments. The isolated PBMCs were then aliquoted for subsequent analyses.

#### 2.3.2 *MTB* Infection and Stimulation

According to the experimental requirements and culture conditions, THP-1-derived macrophages were infected with different *MTB* strains, including *H37Ra*, *BCG*, and GFP-*BCG* at a multiplicity of infection (MOI) of 10, or *H37Rv* at an MOI of 1. In addition, macrophages were stimulated with *H37Rv* lysate obtained from the Youchao Dai Laboratory at final concentrations of 5, 10, 20, 30, or 50 μg/mL. After 6 h of infection with *H37Ra*, *BCG*, or GFP-*BCG* at an MOI of 10, or stimulation with *H37Rv* lysate, the supernatant was removed, and the cells were washed three times with sterile 1× PBS to eliminate extracellular mycobacteria or lysate. The cells were then replenished with complete DMEM and incubated for an additional 18 h.

### 2.4 RNA preparation and Real time PCR

Total RNA was extracted using TRIzol reagent (Invitrogen, USA) in accordance with the manufacturer’s instructions. For each sample, 450 ng of RNA was reverse-transcribed into complementary DNA (cDNA) under the following conditions: 37°C for 15 min, 85°C for 5 s, followed by a 4°C hold. Quantitative real-time PCR was performed using TransStart Tip Green qPCR SuperMix (TransGen, Beijing, China) on a QuantStudio™ 6 Flex Real-Time PCR System (Thermo Fisher Scientific, USA). All reactions were conducted in triplicate, and relative target gene expression was calculated using the 2^-ΔΔCt^ method. The circular RNA detection assay generated a sequence containing the reverse adapter, and its nucleotide composition was determined by sequencing.

### 2.5 Western blot analysis

Protein was extracted by homogenizing 5 × 10^5^ cells in 200 µL of RIPA lysis buffer supplemented with 1% PMSF on ice for 30 min. The lysates were then centrifuged at 12,000 × g for 15 min at 4°C, and the resulting supernatants were collected. Protein concentrations were determined using the Pierce BCA Protein Assay Kit. All measurements were performed in duplicate. Equal amounts of protein were separated by SDS–PAGE and transferred onto PVDF membranes (Millipore, USA). The membranes were blocked with 5% skim milk in TBST for 1 h and incubated overnight at 4°C with the primary monoclonal antibody. After three 10-min washes with TBST, the membranes were incubated with the corresponding HRP-conjugated secondary antibody for 1 h at room temperature. Immunoreactive bands were visualized using an enhanced chemiluminescence detection system and chemiluminescent substrate (Thermo Scientific, USA).

### 2.6 RNase R treatment

A total of 450 ng RNA was treated with 1 U RNase R (Thermo Fisher, USA) in 1× RNase R reaction buffer for 30 min at 37°C. The treated RNA was then reverse-transcribed into cDNA using PrimeScript RT Reagent (TaKaRa, Japan). Quantitative real-time PCR (qRT-PCR) was subsequently performed with TransStart Tip Green qPCR SuperMix (TransGen, Beijing, China) on a QuantStudio™ 6 Flex Real-Time PCR System (Applied Biosystems, Foster City, CA, USA).

### 2.7 RNA pulldown

An amount equivalent to 10⁷ THP-1 cells was used to establish the *MTB* infection cell model. The cells were lysed with IP lysis buffer for 10 min and then centrifuged. The resulting lysates were incubated with 100 nM circ_0074371-or miR-4738-3p-specific probes, or the corresponding control probes, at room temperature for 1 h with rotation at 10 rpm in a DNA rotator. Subsequently, 50 μL of streptavidin magnetic beads (Life Technologies) was added, and the samples were incubated overnight at 10 rpm in the DNA rotator. RNA was then extracted using TRIzol and analyzed by qRT-PCR. The sequences of the probes are provided in Table S4.

### 2.8 Dual luciferase reporter assay

Wild-type and mutant sequences were cloned into the dual-luciferase reporter vector psiCHECK2. The resulting plasmids were co-transfected into 293T cells with miR-4738-3p mimics, inhibitors, or the corresponding negative controls (NC) using Lipofectamine reagent. After 48 h, firefly and Renilla luciferase activities were measured with a luminometer, and the relative light units (RLUs) were calculated. Stronger intermolecular binding resulted in lower Renilla luciferase activity and a reduced Renilla-to-firefly luciferase ratio.

### 2.9 Fluorescence In Situ Hybridization (FISH)

To detect the intracellular localization and colocalization of circ_0074371 and miR-4738-3p, RNA FISH was performed as follows. The biotin-conjugated Circ_0074371 and CY3-conjugated miR-4738-3p probes were synthesised by Shanghai Jima Pharmaceutical Technology Co., Ltd. Briefly,the tuberculosis THP-1 cell model was rinsed in RNase-free PBS and fixed with 4% methanol for 30 min at room temperature.After removal of PBS,the cells were treated with absolute ethanol for 1 min.The circ_0074371 and miR-4738-3p probes were hybridized with the samples overnight at 37°C in the dark.Subsequently, the cells were blocked with 3% bovine serum albumin (BSA) for 30 min and incubated with anti-biotin CY5.5 conjugate and CY3 conjugate for 1 h at 37°C. Following incubation with DAPI for 4 min in the dark, an anti-fluorescence quenching mounting medium was applied. The cells were visualised using laser scanning confocal microscope (Leica, Germany). The probe sequences used are listed in the table S5.

### 2.10 Mycobacteria CFU assay

THP-1-derived macrophages were infected with *H37Ra* at an MOI of 10. 6 hours post-infection, extracellular non-internalised bacilli were removed by washing three times with prewarmed PBS, establishing this time point as 0. Subsequently, the cell supernatant was discarded, and the infected cells were lysed in 1 mL sterile water containing 0.03% sodium dodecyl sulfate (SDS) for 20 mins. A 10-fold serial dilution was conducted for quantitative culture analysis. Aliquots of 100 µL were plated in triplicate on Middlebrook 7H11 agar supplemented with 10% oleic acid-albumin-dextrose-catalase (OADC) enrichment and incubated for 2–3 weeks until colonies reached adequate size for enumeration. The viability of Mycobacteria was assessed by counting colony-forming units (CFU).

### 2.11 Flow cytometry analysis

The fluorescence detected in this study originated from the intrinsic fluorescence of the bacterial or cell strains. GFP-*BCG* fluorescent bacteria were analyzed using a BD FACS-Verse flow cytometer, and FITC fluorescence in cell lines with overexpression of circ_0074371 was acquired through sorting with a CytoFLEX flow cytometer.

### 2.12 ELISA detection

Plasma or culture medium levels of IL-10 and TNF-α were quantified using commercially available ELISA kits, adhering to the manufacturer’s instructions. The IL-10 and TNF-α ELISA kits utilized were supplied by eBioscience (product BMS215/2, BMS223/4; eBioscience, San Diego, CA).

### 2.13 In vitro stimulation experiment

PBMCs were maintained in RPMI 1640 medium supplemented with 10% fetal bovine serum, 100 units/ml penicillin, and 100 μg/ml streptomycin at 37°C in a humidified incubator with 5% CO2. The cells were seeded in 24-well microplates at a concentration of 1×10^6^ cells/well, followed by adding living *BCG* and lipopolysaccharide (LPS) respectively for 6, 12 and 24 hrs. *BCG* was washed and suspended with PBS (0.05% Tween 80). The concentration of *BCG* is MOI=1:20. 1 μg/ml lipopolysaccharide (LPS) was used as a positive control and PBS (0.05% Tween 80) was used as a negative control. After culture for an indicated period, cell culture supernatants were collected, filtered through a 0.22 μm filter, and stored at-80°C until analysis.

### 2.14 Macrophage polarization detection

Macrophage polarization is characterized by detecting polarization-related factors. For M1-type polarization, the selected factors are TNF-α, Nos2, and IL-12p40; for M2-type polarization, the selected factors are IL-10, Chi3l1, and Arg1. The expression levels of mRNA were detected using Real-time qPCR, and the expression levels of polarization factor proteins were detected using Western blotting.

### 2.15 Statistical analysis

Normality test was first performed to decide whether the data were in normal distribution. Data were shown as Mean ± SEM (standard error of the mean, SEM). Student’s *t*-test or *ANOVA* (analysis of variance) were employed to compare the differences of measured data, and *Pearson* correlation was used to measure the degree of dependency between variables by the GraphPad Prism version 5.0 software (GraphPad Software Inc., San Diego, CA, USA). A P value of less than 0.05 was considered as statistically significant.

## Results

### Marked Downregulation of Circ_0074371 in Patients with Active Tuberculosis Supports Its Potential as a Novel Diagnostic Biomarker

Previously, using circRNA-seq high-throughput sequencing, we identified numerous differentially expressed circRNAs in peripheral blood mononuclear cells (PBMCs) from patients with active pulmonary tuberculosis (ATB) compared with healthy controls (HCs) (Mol Immunol. 2017). In the present study, we selected several circRNAs with relatively high differential expression levels for further investigation. Among these, circ_0074371 was significantly downregulated in patients with ATB compared with HCs (Figure 1A). To further validate this finding, peripheral blood samples were collected from 32 patients with ATB and 37 HCs. RT-qPCR analysis confirmed that circ_0074371 expression in the PBMCs of patients with ATB was significantly lower than that in the HC group (Figure 1B). In addition, circ_0074371 expression was significantly increased after one month of standardized anti-tuberculosis treatment (M1) compared with baseline levels before treatment (M0) (Figure 1C). Furthermore, the area under the curve (AUC) for circ_0074371 in the diagnosis of ATB was 0.8654 (P < 0.0001; Figure 1D), suggesting that circ_0074371 may serve as a diagnostic biomarker for ATB with good diagnostic performance.

**Figure 1:**
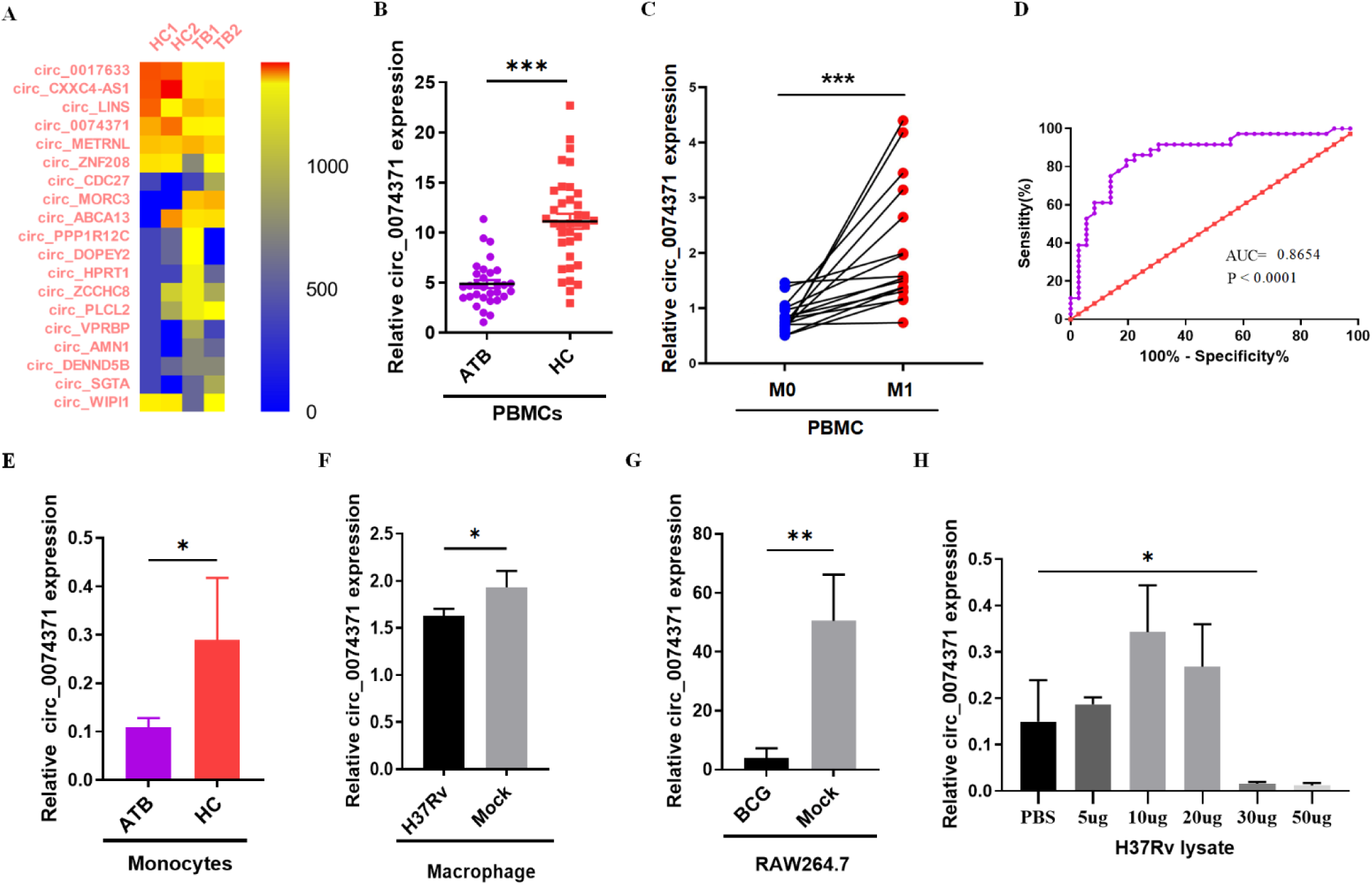
circ_0074371 is significantly downregulated in patients with active tuberculosis and may serve as a novel biomarker for its diagnosis. (A) Heatmap showing circular RNAs that were differentially expressed in PBMCs from patients with active TB compared with healthy controls (HCs). (B) circ_0074371 expression was further assessed by qRT-PCR in PBMCs from 31 patients with active TB and 36 HCs. Actin was used as the housekeeping gene to normalize circRNA expression. Data are presented as the mean ± SEM. ***P < 0.001 vs HC. (C) circ_0074371 expression was evaluated in PBMCs from 16 patients with active TB before (pre-T) and after (post-T) 1 month of individualized anti-TB treatment with isoniazid, rifampicin, pyrazinamide, and ethambutol. Data are presented as the mean ± SEM. ***P < 0.001 vs pre-T. (D) The diagnostic value of circ_0074371 for active TB was evaluated using receiver operating characteristic (ROC) curve analysis. (E) circ_0074371 expression was further examined by qRT-PCR in peripheral blood monocytes from 5 patients with active TB and 5 HCs. Peripheral blood monocytes were isolated from PBMCs by immunomagnetic positive selection. *P < 0.05 vs HC. (F) THP-1 macrophages were infected with *H37Rv* at a multiplicity of infection (MOI) of 1, after which circ_0074371 expression was measured by qRT-PCR. Data were obtained from three independent experiments. *P < 0.05 vs Mock. (G) THP-1 macrophages were infected with *BCG* at an MOI of 10, and circ_0074371 expression was subsequently measured by qRT-PCR. Data were obtained from three independent experiments. **P < 0.01 vs Mock. (H) THP-1 macrophages were treated with different concentrations of *H37Rv* lysate (0, 5, 10, 20, 30, and 50 μg), and circ_0074371 expression was measured by qRT-PCR. Data were obtained from three independent experiments. *P < 0.05 vs Mock. ATB, active tuberculosis; HC, health control; PBMCs, peripheral blood mononuclear cells; qRT-PCR, quantitative real-time polymerase chain reaction; ROC, reactive operating characteristic curve; *H37Rv*, Mycobacterium tuberculosis *H37Rv*; *BCG*, Bacillus Calmette-Guérin; MOI, multiplicity of infection.

Because PBMCs comprise a heterogeneous cell population, we further investigated circ_0074371 expression in isolated cell populations from HC and ATB patients. Monocytes were isolated from PBMCs using magnetic bead-based sorting. RT-qPCR analysis revealed that circ_0074371 expression was significantly lower in monocytes from ATB patients than in those from the HC group (Figure 1E). Consistently, in an in vitro infection model, circ_0074371 expression was significantly reduced in THP-1-derived macrophages infected with *H37Rv* or *BCG* compared with the corresponding control group (Figures 1F and 1G). Moreover, treatment of THP-1 cells with increasing concentrations of *H37Rv* lysate revealed a significant downregulation of circ_0074371 expression at a lysate concentration of 30 μg (Figure 1H), This pattern was consistent with that observed following whole-bacterial infection; therefore, *H37Rv* lysate was used in subsequent experiments to stimulate the cells. Taken together, these findings indicate that circ_0074371 is significantly downregulated in patients with ATB and may serve as a novel biomarker for the diagnosis of active pulmonary tuberculosis.

### Circ_0074371 expression pattern in *MTB*-infected macrophages

Next, we investigated the subcellular localization of circ_0074371 in macrophages. Bioinformatic analysis revealed that circ_0074371 originates from the host gene Rho GTPase-activating protein 26 (ARHGAP26) and is generated by back-splicing of exons 15, 16, and 17, resulting in a full-length transcript of 253 bp (Figure 2A). We then designed primers spanning the back-splicing junction for RT-qPCR and performed Sanger sequencing of the amplified products. The sequencing results confirmed that the products were identical to the back-spliced junction sequence of circ_0074371 (Figure 2B). To further verify the circular RNA characteristics of circ_0074371, total RNA was treated with RNase R. RT-qPCR analysis showed that circ_0074371 was significantly more resistant to RNase R digestion than linear actin RNA (Figures 2C, 2D). In parallel, THP-1 cells were treated with siRNA targeting circ_0074371. Agarose gel electrophoresis of the RT-qPCR products from both the siRNA-treated and RNase R-treated groups showed bands at the expected positions, further confirming that circ_0074371 exists as a stable circular transcript. Moreover, RNA fluorescence in situ hybridization (FISH) and nuclear–cytoplasmic fractionation assays demonstrated that circ_0074371 was predominantly localized in the nucleus. Following *Mycobacterium tuberculosis* infection, circ_0074371 expression decreased markedly in the nucleus but increased in the cytoplasm.

**Figure 2:**
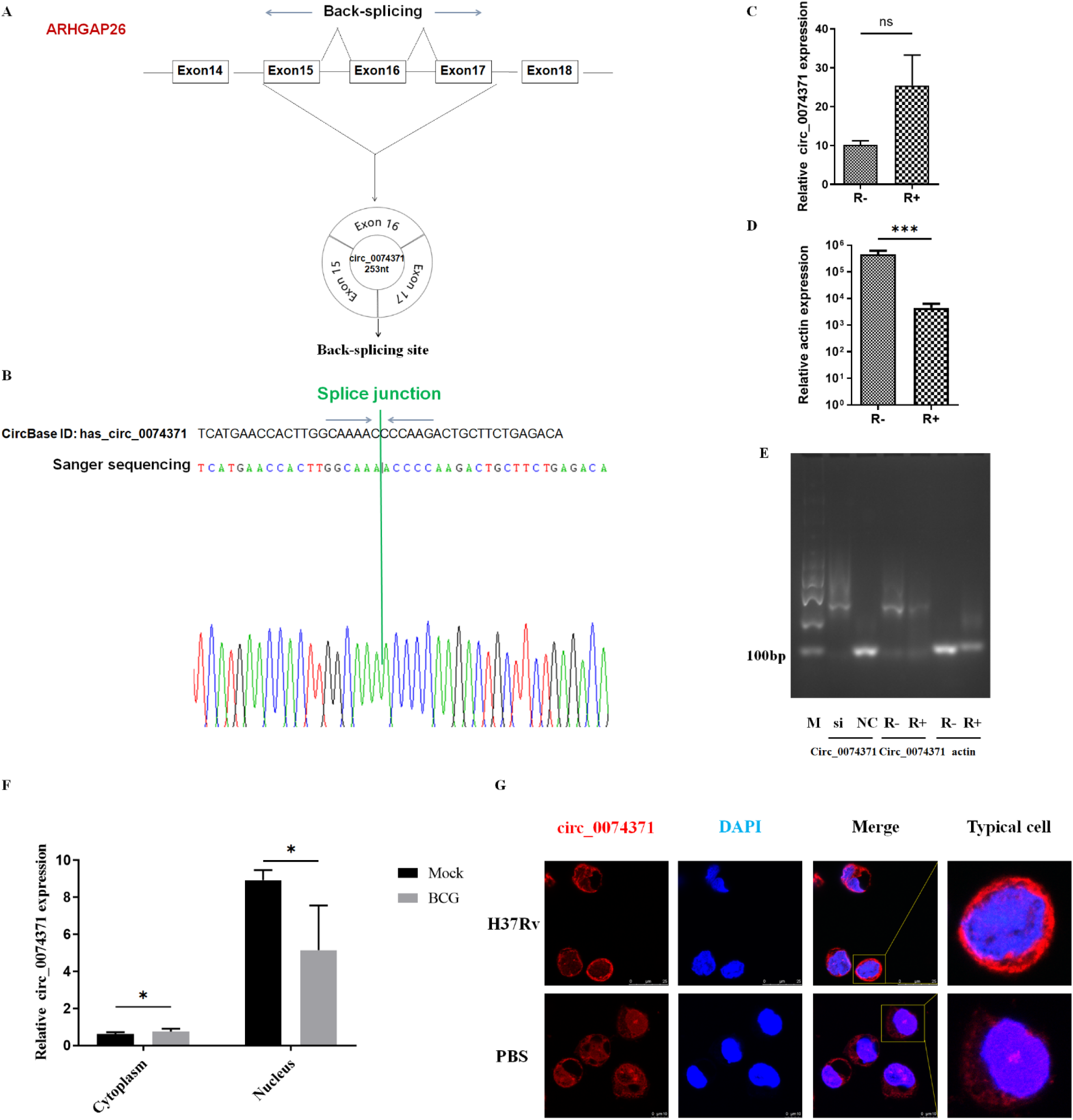
Circ_0074371 expression pattern in *MTB*-infected macrophages. (A) Schematic illustration of the circularization of exons 15–17 of the ARHGAP26 gene to form circ_0074371. (B) The splice junction sequence of circ_0074371 recorded in CircBase (circular RNA database; upper panel) was consistent with the sequence obtained by Sanger sequencing of qRT-PCR products (lower panel). (C–D) Nucleic acids from THP-1 macrophages were treated with RNase R (R+) or left untreated (R−), followed by qRT-PCR analysis of circ_0074371 and linear Actin mRNA expression. Data were obtained from three independent experiments and are presented as the mean ± SEM. ***P < 0.001 versus R+. (E) Following transfection with si-circ_0074371 or si-NC and treatment with RNase R (R+) or no treatment (R−), qRT-PCR products were analyzed by agarose gel electrophoresis. (F) After cytoplasmic and nuclear RNA separation, the subcellular distribution of circ_0074371 in THP-1 macrophages infected with *BCG* at an MOI of 10 for 24 h was assessed by qRT-PCR. Data were obtained from three independent experiments. *P < 0.05 versus the mock group. (G) THP-1 macrophages were stimulated with *H37Rv* lysate (30 μg/mL) for 24 h and then incubated overnight at 37°C with a biotin-conjugated circ_0074371 probe. A representative immunofluorescence confocal image showing the subcellular localization of circ_0074371 is presented. qRT-PCR, quantitative real-time polymerase chain reaction; *BCG*, Bacillus Calmette-Guerin; MOI, multiplicity of infection; FISH, fluorescence in situ hybridisation.

### Circ_0074371 enhances the antimicrobial activity of macrophages against intracellular Mycobacterium tuberculosis

The marked downregulation of circ_0074371 in patients with active pulmonary tuberculosis suggests that it may contribute to the pathogenesis of active tuberculosis (ATB). Because macrophages are the primary host cells targeted by M. tuberculosis and the intracellular survival of the bacterium within macrophages is closely associated with the development of ATB, we next examined whether the altered expression of circ_0074371 directly affects intracellular M. tuberculosis survival. Macrophages were infected with genetically engineered GFP-expressing Bacille Calmette– Guérin (GFP-*BCG*) while simultaneously undergoing circ_0074371 knockdown. Compared with the control group, the siRNA-treated group exhibited a higher proportion of GFP-positive macrophages and increased mean fluorescence intensity (Figures 3A–C). RT-qPCR confirmed that the siRNA effectively reduced circ_0074371 expression (Figures 3D, 3E). We subsequently generated a circ_0074371 overexpression vector using a lentiviral system and transduced THP-1 cells to establish a stable circ_0074371-overexpressing cell line. Fluorescence microscopy showed strong and consistent expression of the vector-derived fluorescence, and flow cytometric sorting yielded a purified cell population with an overexpression positivity rate of 98% (Supplementary Figure S2A, 2B). Importantly, RT-qPCR analysis showed that the expression of the linear ARHGAP26 transcript was not significantly altered in the circ_0074371-overexpressing cell line (Figures 3F, 3G). To further determine the effect of circ_0074371 on intracellular M. tuberculosis survival, we performed colony-forming unit (CFU) assays using GFP-positive cells, as confirmed by flow cytometry, isolated from the circ_0074371-overexpressing cell line (Figures 3H, 3I). Following infection with *H37Ra* at a multiplicity of infection of 10 for 24 h, CFU counts were significantly lower in circ_0074371-overexpressing THP-1 macrophages than in LV-NC THP-1 macrophages (Figure 3J). These findings indicate that circ_0074371 overexpression suppresses the intracellular growth of M. tuberculosis in macrophages.

**Figure 3:**
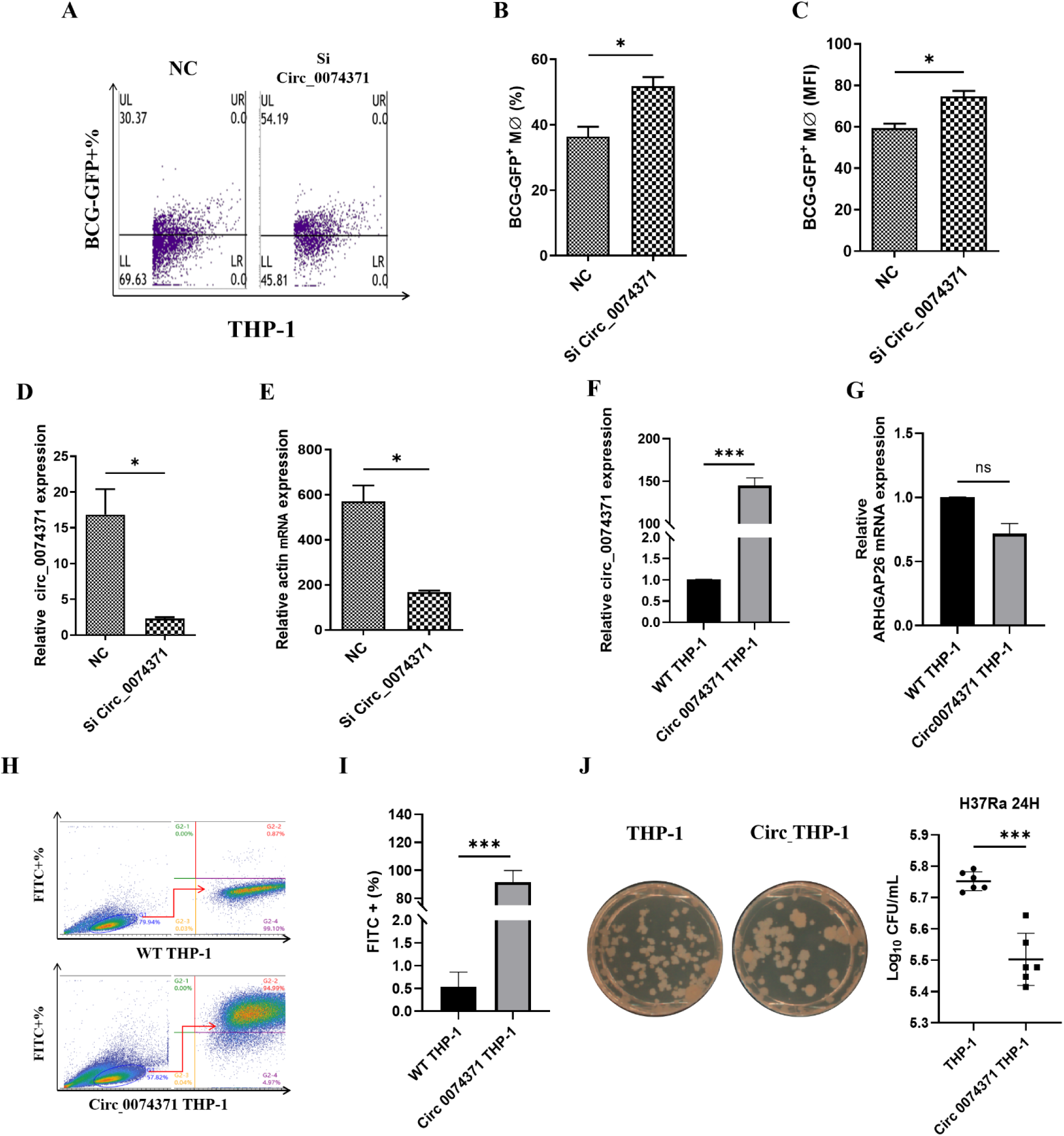
Circ_0074371 enhances the antimicrobial activity of macrophages against intracellular Mycobacterium tuberculosis. (A) Representative flow cytometry histograms showing the percentage of macrophages expressing fluorescent *BCG* (*BCG*-GFP⁺ macrophages) after transfection with siRNA targeting circ_0074371. (B) Quantification of the percentage of *BCG*-GFP⁺ macrophages. Data were obtained from three independent experiments. *P < 0.05 vs. NC. (C) Quantification of the mean fluorescence intensity of *BCG*-GFP⁺ macrophages. Data are presented as the mean ± SEM. *P < 0.05 vs. NC. (D) Relative expression levels of circ_0074371. Data were obtained from three independent experiments. *P < 0.05 vs. NC. (E) Relative expression levels of actin mRNA, used as a positive control. Data were obtained from three independent experiments. *P < 0.05 vs. NC. Si, siRNA targeting circ_0074371; NC, negative control siRNA for circ_0074371. (F) Relative expression levels of circ_0074371 in the constructed circ_0074371-overexpressing cell line. Data were obtained from three independent experiments. ***P < 0.001 vs. wild-type (WT) THP-1 macrophages. (G) Relative expression levels of circ_ARHGAP26 mRNA in the constructed circ_0074371-overexpressing cell line. Data were obtained from three independent experiments. ***P < 0.001 vs. WT THP-1 macrophages. (H) Representative flow cytometry histograms showing the percentage of FITC-labeled THP-1 cells in WT THP-1 and circ_0074371-overexpressing THP-1 cell lines. (I) Quantification of the percentage of FITC⁺ THP-1 cells. Data were obtained from three independent experiments. ***P < 0.001 vs. WT THP-1 macrophages. (J) Schematic illustration of the bactericidal activity assay and statistical analysis of colony counts in WT THP-1 and circ_0074371-overexpressing THP-1 macrophages infected with *H37Ra* for 24 h at an MOI of 10. Data were obtained from three independent experiments. ***P < 0.001.

### Circ_0074371 promotes M1 polarization via AMPK/mTOR signaling pathway in *MTB* infected macrophages

Given the effect of Circ_0074371 on the intracellular survival of mycobacteria, we next investigated whether it restricts mycobacterial growth by modulating macrophage function. We first assessed whether macrophage autophagy contributes to the survival of intracellular tuberculosis bacteria by measuring the autophagy marker LC3B I/II using Western blotting. Circ_0074371 overexpression had no detectable effect on autophagy (Figure S4). Because macrophage polarization is important for host defense against *MTB*, we then examined whether Circ_0074371 affects macrophage bactericidal activity by altering their polarization state. RT-qPCR analysis of polarization markers following *BCG* infection showed that, compared with the control group, Circ_0074371 overexpression significantly increased the expression of M1-associated factors and decreased that of M2-associated factors (Figure 4A). We further evaluated polarization-related factors in *MTB*-infected macrophages using Western blotting and ELISA. TNF-α and IL-10 were selected as representative cytokines of the M1 and M2 phenotypes, respectively. Circ_0074371 overexpression significantly increased TNF-α expression (Figure 4B–D) while reducing IL-10 expression (Figure 4E–G). Together, these findings indicate that Circ_0074371 overexpression promotes macrophage polarization toward the M1 phenotype. Our previous findings demonstrated that Circ_0074371 promotes M1 polarization in mycobacteria-infected macrophages. Given the important role of the AMPK/mTOR signaling pathway in macrophage polarization, we further examined the phosphorylation status of AMPK and mTOR by Western blotting. Circ_0074371 overexpression significantly reduced AMPK phosphorylation (Figure 4H–K) while enhancing mTOR activation (Figure 4L–O). These findings suggest that Circ_0074371 promotes M1 macrophage polarization by suppressing AMPK phosphorylation and activating mTOR. Collectively, these data demonstrate that Circ_0074371 regulates the polarization of *MTB*-infected macrophages.

**Figure 4:**
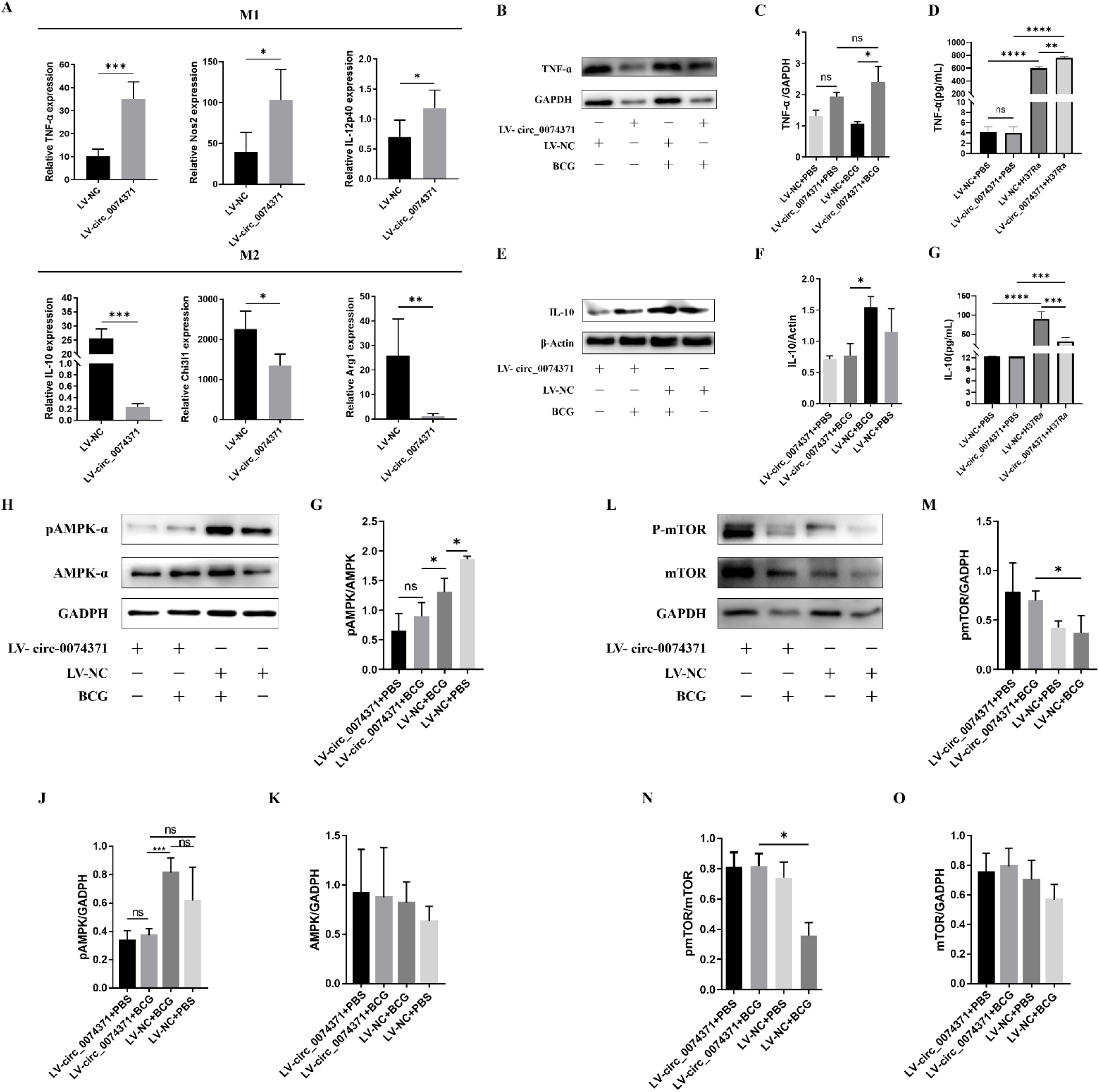
Circ_0074371 promotes M1 polarization via AMPK/mTOR signaling pathway in *MTB* infected macrophages. (A) THP-1 macrophages were transduced with a lentiviral vector overexpressing circ_0074371 (LV-circ_0074371) or the corresponding negative control vector (LV-NC), then infected with *BCG* for 24 h. The expression of M1-and M2-associated genes was subsequently analyzed by RT-PCR. (B, C) THP-1 macrophages were infected with *BCG* at a multiplicity of infection (MOI) of 10 for 24 h. Representative Western blots and quantitative analysis of TNF-α protein expression in THP-1 macrophages are shown. (D) ELISA was used to determine TNF-α levels in THP-1 macrophages. (E, F) Representative Western blots and quantitative analysis of IL-10 protein expression in THP-1 macrophages are shown. (G) ELISA was performed to measure IL-10 levels in THP-1 macrophages. (H–K) Representative Western blots and quantitative analysis of phosphorylated AMPK-α and total AMPK-α protein expression in THP-1 macrophages are shown. (L–O) Representative Western blots and quantitative analysis of phosphorylated mTOR and total mTOR protein expression in THP-1 macrophages are shown. All data were obtained from three independent experiments and are presented as the mean ± SEM. *P < 0.05, **P < 0.01, and ***P < 0.001 versus the LV-NC group.

### circ_0074371 Interacts with miR-4738-3p, Which Is Upregulated During Mycobacterial Infection

The principal mechanism through which circRNAs exert their biological functions is by binding to miRNAs and subsequently regulating the expression of miRNA target proteins. To elucidate how Circ_0074371 regulates mycobacterial growth and macrophage polarization, we used the public databases miRWalk, DIANA, TargetScan, miRTarBase, and miRDB to predict its potential target miRNAs. A cross-database analysis identified miR-4738-3p as a candidate miRNA capable of binding to Circ_0074371 (Figure 5A). We further found that miR-4738-3p expression was significantly higher in PBMCs from patients with ATB than in those from healthy controls (Figure 5B). Similarly, miR-4738-3p expression was upregulated in THP-1 macrophages infected with *BCG* in vitro, whereas Circ_0074371 knockdown further increased miR-4738-3p expression (Figure 5C). In addition, FISH was performed to determine the subcellular localization of miR-4738-3p. In THP-1 macrophages treated with *H37Rv* lysate, the cytoplasmic red fluorescence corresponding to miR-4738-3p was markedly stronger than that observed in the control group (Figure 5D). These findings suggest that miR-4738-3p is closely associated with mycobacterial infection.

**Figure 5:**
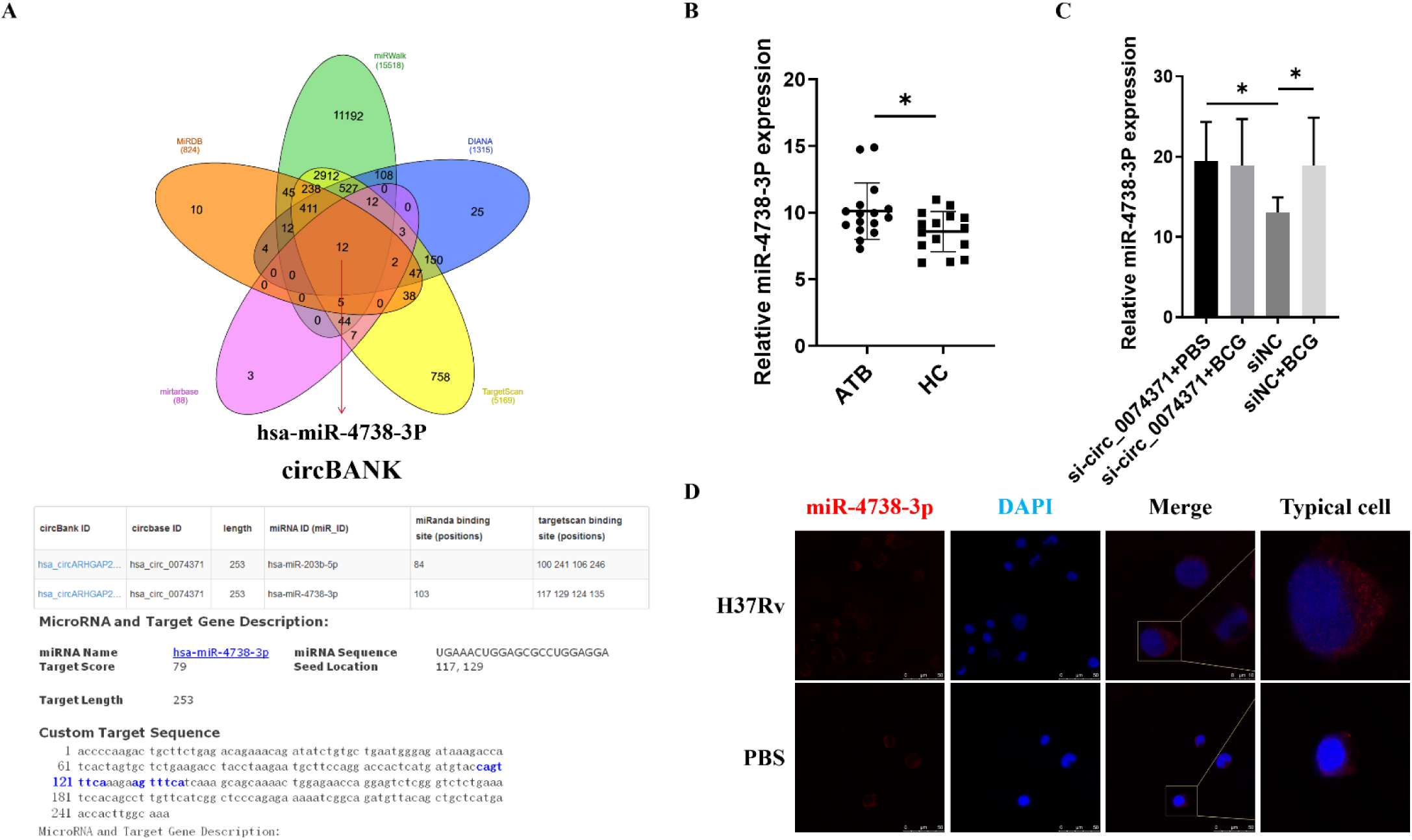
Interaction Between circ_0074371 and miR-4738-3p, Which Is Upregulated During Mycobacterial Infection. (A) Using the circRNA prediction databases miRWalk, DIANA, TargetScan, miRTarBase, and miRDB, we identified miR-4738-3p as a potential target of circ_0074371. The schematic diagram illustrates the predicted binding sites and the corresponding mutation sites between circ_0074371 and miR-4738-3p. (B) The relative expression level of miR-4738-3p in human PBMCs from 16 patients with active TB and 15 healthy controls (HCs) was determined by qRT-PCR. Actin was used as the housekeeping gene for normalization. Data are presented as the mean ± SEM. *P < 0.05 versus HC. (C) The relative expression level of miR-4738-3p was measured by qRT-PCR in *BCG*-infected THP-1 macrophages following transfection with siRNA targeting circ_0074371. The data were obtained from three independent experiments and are presented as the mean ± SEM. *P < 0.05 versus siNC. (D) FISH was performed to examine the cytoplasmic localization of miR-4738-3p in *H37Rv*-infected THP-1 macrophages (×400). *H37Rv* infection increased the expression of miR-4738-3p. FISH, fluorescence in situ hybridization.

### circ_0074371 Directly Binds to miR-4738-3p

circ_0074371 and miR-4738-3p were significantly associated with the regulation of macrophage function, suggesting a potential direct interaction between these two molecules. To investigate this possibility, bioinformatics analysis was first performed to predict the putative binding sites between circ_0074371 and miR-4738-3p (Figure 6A). Dual-luciferase reporter vectors containing either the wild-type circ_0074371 sequence (circ_0074371-WT) or a sequence with a mutated miR-4738-3p binding site (circ_0074371-Mut) were constructed and cotransfected with a miR-4738-3p mimic or inhibitor. No significant differences in luciferase activity were observed in the empty-vector group under any treatment condition (Figure 6B). In the WT group, the luciferase activity ratio was significantly reduced by the miR-4738-3p mimic and increased by the miR-4738-3p inhibitor (Figure 6C). However, neither treatment significantly altered the luciferase activity ratio in the Mut group (Figure 6D).

**Figure 6:**
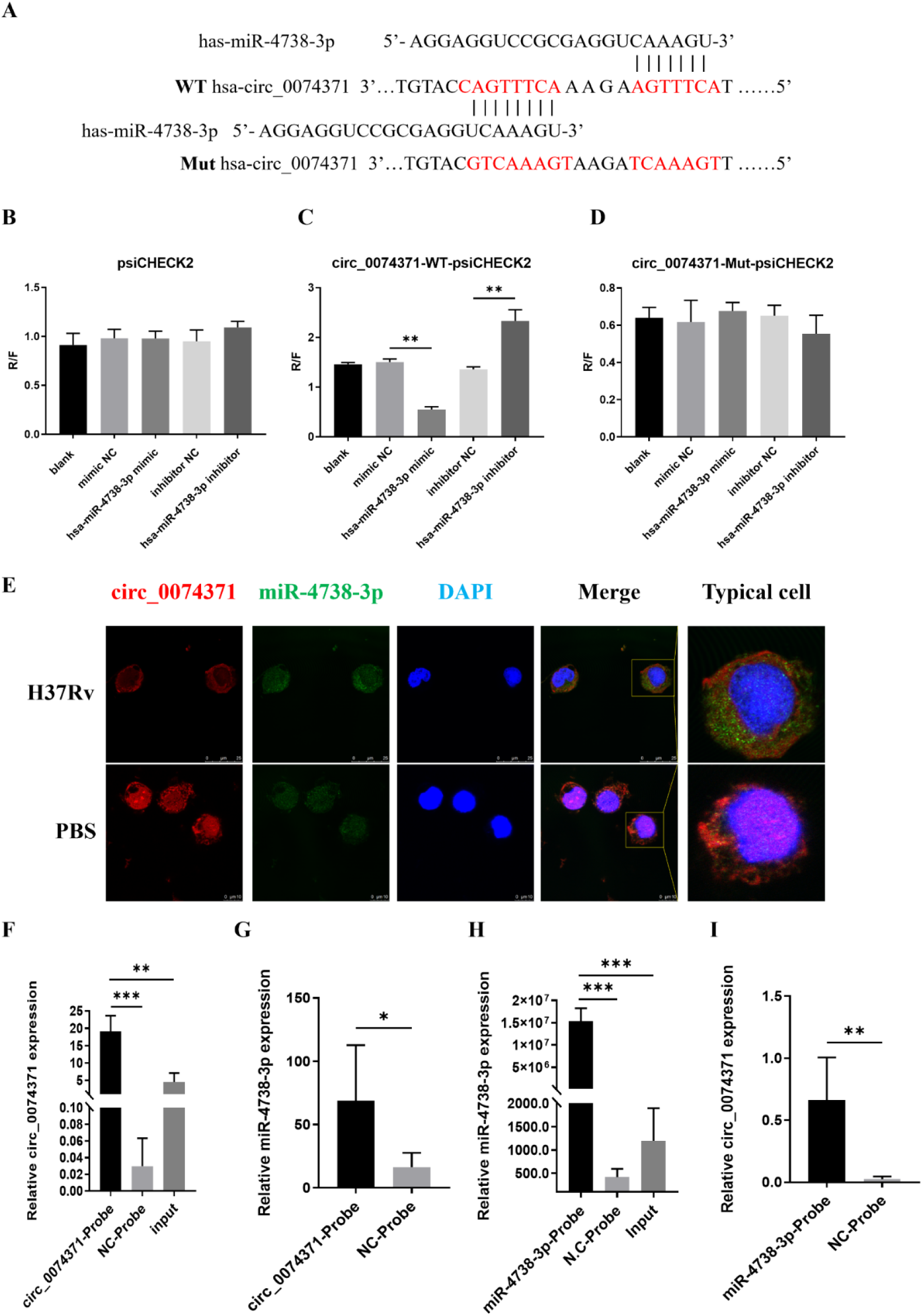
Direct Binding of circ_0074371 with miR-4738-3p. (A) Schematic illustration of the predicted binding site between circ_0074371 3′UTR and miR-4738-3p. (B–D) Relative luciferase activity was assessed in HEK293T cells co-transfected with miR-4738-3p mimics or inhibitors and psiCHECK2, circ_0074371-WT-psiCHECK2, or circ_0074371-MUT-psiCHECK2. Data were obtained from three independent experiments and are presented as the mean ± SEM. **P < 0.01 versus NC. (E) Co-localization of miR-4738-3p and circ_0074371 in *BCG*-infected THP-1 macrophages was examined by FISH (×400). (F, G) A circRNA pull-down assay was performed in *BCG*-infected THP-1 macrophages using biotin-labeled circ_0074371 to evaluate its interaction with miR-4738-3p. The enrichment of circ_0074371 and miR-4738-3p was subsequently quantified by qRT-PCR. Data were obtained from three independent experiments and are presented as the mean ± SEM. *P < 0.05, **P < 0.01, and ***P < 0.001 versus the control probe. (H, I) A miRNA pull-down assay was performed in *BCG*-infected THP-1 macrophages using biotin-labeled miR-4738 probes, followed by qRT-PCR analysis to determine the enrichment of miR-4738-3p and circ_0074371. Data were obtained from three independent experiments and are presented as the mean ± SEM. **P < 0.01 and ***P < 0.001 versus the control probe.

The colocalization of circ_0074371 and miR-4738-3p was further examined using RNA fluorescence in situ hybridization (RNA FISH). Following treatment with *H37Rv* lysate, circ_0074371 expression was reduced, whereas the cytoplasmic green fluorescence signal corresponding to miR-4738-3p was markedly increased compared with that in the control group. In addition, circ_0074371 and miR-4738-3p were observed to colocalize in the cytoplasm (Figure 6E). RNA pull-down assays showed that, in *BCG*-infected macrophages, the circ_0074371 and miR-4738-3p probes pulled down significantly higher levels of miR-4738-3p and circ_0074371, respectively, than the corresponding control probes (Figure 6F–I). These findings confirm a molecular interaction between circ_0074371 and miR-4738-3p. Collectively, the results demonstrate that circ_0074371 directly targets and binds miR-4738-3p, indicating a significant regulatory relationship between the two molecules.

### Circ_0074371 inhibits AMPK activation to regulate M1 macrophage polarization via target miR-4738-3p

To further investigate the role of miR-4738-3p in mediating the effects of circ_0074371 on macrophage polarization during Mycobacterium tuberculosis infection, an in vitro macrophage model was established by liposome-mediated transfection with a miR-4738-3p mimic or inhibitor. Western blot analysis showed that the miR-4738-3p mimic increased the level of phosphorylated AMPK (Figure 7A–B), whereas the miR-4738-3p inhibitor acted synergistically with circ_0074371 to suppress AMPK phosphorylation (Figure 7C–D). Moreover, following treatment with *H37Rv* lysate, the miR-4738-3p mimic enhanced AMPK phosphorylation and reduced TNF-α expression (Figure 7E–G). Conversely, the miR-4738-3p inhibitor synergized with circ_0074371 to inhibit AMPK phosphorylation and increase TNF-α expression (Figure 7H–J). Taken together, these findings indicate that the circ_0074371/miR-4738-3p axis promotes M1 macrophage polarization in response to mycobacterial infection by regulating the AMPK signaling pathway.

**Figure 7:**
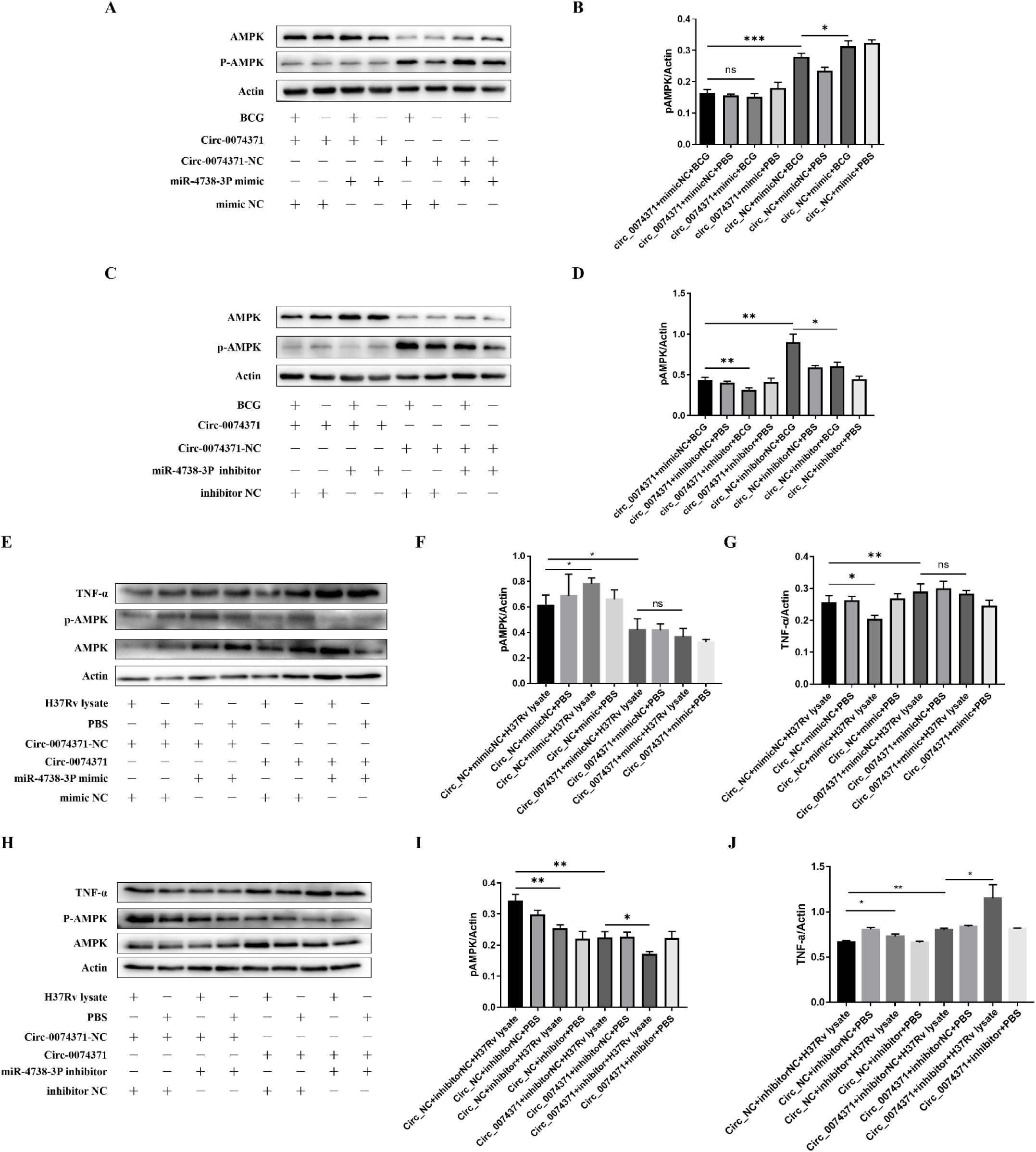
Circ_0074371 inhibits AMPK activation to regulate M1 macrophage polarization via miR-4738-3p. (A–D) THP-1 macrophages were transduced with a lentiviral vector overexpressing circ_0074371 (LV-circ_0074371) or the corresponding negative-control vector (LV-NC), and were simultaneously treated with miR-4738-3p mimics or mimic negative control (mimics NC), or with a miR-4738-3p inhibitor or inhibitor negative control (inhibitor NC). Representative Western blot images showing the expression of phosphorylated AMPK-α and total AMPK-α in macrophages infected with *BCG* for 24 h are presented. The results were obtained from three independent experiments and are expressed as the mean ± SEM. ***P < 0.001, **P < 0.01, and *P < 0.05. (E–J) THP-1 macrophages were also treated with *H37Rv* lysate. Representative Western blot images showing the expression of TNF-α, phosphorylated AMPK, and total AMPK are presented, with results consistent with those observed following *BCG* infection. The results were obtained from three independent experiments and are expressed as the mean ± SEM. **P < 0.01 and *P < 0.05.

## Discussion

In this study, we identified a novel competing endogenous RNA (ceRNA) regulatory axis that links circRNA activity with macrophage polarization and the intracellular clearance of mycobacteria during tuberculosis (TB). The polarization state of pulmonary macrophages in patients with TB is closely associated with host-mediated control of mycobacterial infection^[14,37]^, In vitro studies have shown that promoting M1 macrophage polarization suppresses the intracellular growth of *Mycobacterium tuberculosis*(*MTB*)^[38,39]^, whereas M2 polarization facilitates bacterial persistence and infection^[40]^, Thus, modulating the polarization phenotype of pulmonary macrophages may represent an effective strategy for TB control. Circular RNAs (circRNAs) are a class of highly stable noncoding RNAs that have attracted considerable attention because of their involvement in a wide range of diseases, including cancer, cardiovascular disease, and autoimmune disorders ^[41]^, Their remarkable stability and evolutionary conservation, together with their potential clinical applicability, make circRNAs promising candidates for diagnostic and therapeutic development^[42]^. However, only a limited number of circRNAs associated with *MTB* infection have been well characterized. In our previous study, we identified multiple circRNAs associated with tuberculosis through circRNA transcriptional profiling of peripheral blood mononuclear cells (PBMCs) from patients with active tuberculosis (ATB)^[25]^. In the present study, we identified circ_0074371 as a critical regulator of innate anti-TB immune responses and intracellular *MTB* survival. Mechanistically, circ_0074371 suppresses intracellular *MTB* survival by acting as a competing endogenous RNA to sponge miR-4738-3p, thereby promoting M1 macrophage polarization through inhibition of the AMPK signaling pathway. Analysis of clinical samples revealed that the reduced expression of circ_0074371 in patients with ATB had significant diagnostic value. Functionally, circ_0074371 exhibited substantial biological activity, significantly promoted M1 macrophage polarization, and strongly inhibited AMPK phosphorylation. Collectively, these findings suggest that circ_0074371 may represent a promising immunomodulator for the treatment of tuberculosis(Fig 8).

**Figure 8:**
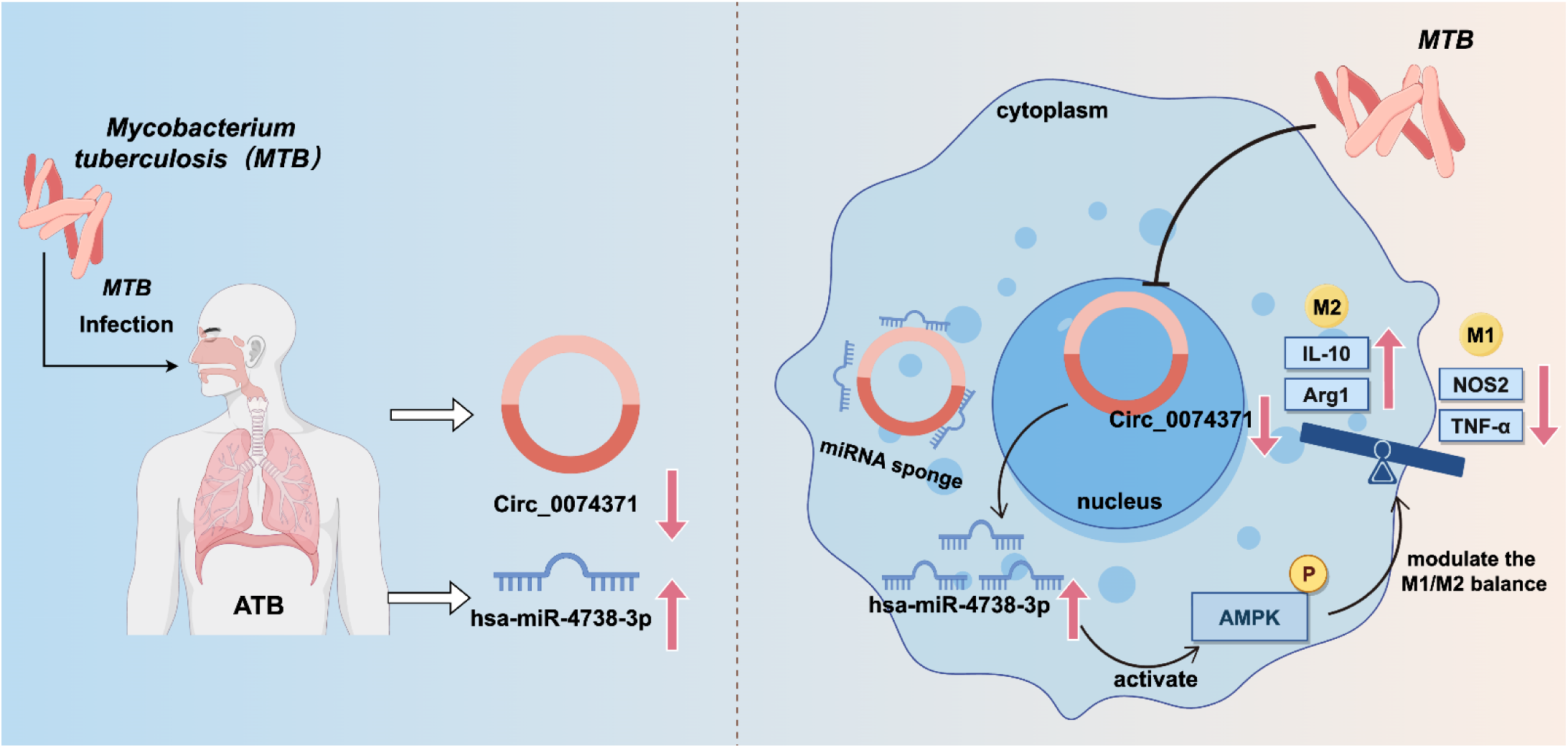
Tuberculosis Patients and a Model Illustrating the Role of the circ_0074371/miR-4738-3p Axis in *MTB*– Macrophage Interactions. circ_0074371 is downregulated during *MTB* infection. By acting as a sponge for miR-4738-3p, it increases miR-4738-3p levels, promotes AMPK phosphorylation, induces M2 macrophage polarization, and favors the development of active tuberculosis. Overexpression of circ_0074371 or inhibition of miR-4738-3p can reverse M1 macrophage polarization, suppress AMPK activity, and enhance macrophage-mediated killing of intracellular mycobacteria.

Although circ_0074371 has been implicated in disease progression in sepsis^[43,44]^, its specific biological functions remain unclear, and its role in patients with tuberculosis has not previously been reported. In the present study, analyses of host cells from clinical patients, together with in vitro infection experiments using tubercle bacilli, demonstrated that marked downregulation of circ_0074371 is closely associated with tuberculosis infection and disease development. Furthermore, cell models with circ_0074371 overexpression or knockdown were established, confirming that this circRNA significantly affects the intracellular survival of tubercle bacilli. Although its effects on pulmonary tuberculosis infection could not be further validated in vivo, the potential role of circ_0074371 in disease progression, as well as its possible utility as a clinical diagnostic biomarker and therapeutic target, should not be overlooked. Relatively few studies have examined the regulation of macrophage polarization by circRNAs in tuberculosis models. Nevertheless, circRNAs capable of modulating macrophage polarization have been identified in several other disease models^[45–47]^, indicating that circRNAs may reshape macrophage polarization phenotypes. In this study, we demonstrated that circ_0074371 promotes M1 macrophage polarization at the transcriptional, protein-expression, and cellular cytokine-secretion levels. Although its effects on pulmonary macrophage polarization have not yet been confirmed in vivo, our findings suggest that circ_0074371 may contribute to remodeling the polarization phenotype of macrophages in the microcirculation.

M1-polarized macrophages can eliminate intracellular *MTB* through multiple mechanisms. As a chronic infectious disease, tuberculosis induces extensive metabolic reprogramming in the host^[48–50]^, thereby influencing macrophage polarization and functional phenotypes. Accordingly, reprogramming macrophage polarization represents a critical strategy for reversing disease progression. M1 polarization promotes the expression of inducible nitric oxide synthase (iNOS, also known as NOS2), which uses arginine as a substrate to generate high levels of nitric oxide (NO). NO can directly target intracellular pathogens that have escaped from the phagosome^[51]^; In addition, M1-polarized macrophages produce abundant reactive oxygen species (ROS), which disrupt the nucleic acid and protein structures of pathogens and thereby restrict the intracellular growth of *MTB*^[52]^; In vivo, M1 macrophages also secrete pro-inflammatory cytokines, including IL-1β, TNF-α, and IL-6, to recruit antigen-specific T cells to the sites of infection^[53,54]^, These responses enhance pathogen-specific immunity and facilitate the elimination of infected target cells. Although the present study examined the effects of macrophage polarization on cytokine production, the intracellular molecular mechanisms by which macrophage polarization contributes to the killing of *MTB* remain unclear. In future studies, using both cellular and animal models, we will investigate the mechanisms underlying the bactericidal activity of this circRNA against *MTB*, with particular emphasis on whether it acts by modulating specific immune responses or by activating intracellular bactericidal pathways.

The principal mechanism through which circ_0074371 exerts its effects appears to involve its direct interaction with miR-4738-3p, thereby promoting macrophage polarization. Bioinformatic analysis initially identified a potential structural basis for the interaction between these two molecules. Subsequent analysis of clinical samples showed that miR-4738-3p was significantly upregulated in patients with ATB. In vitro, miR-4738-3p expression was markedly suppressed by circ_0074371 and was inversely correlated with circ_0074371 expression, which was consistent with the expected functional relationship between the two molecules. Cellular and molecular experiments further confirmed their direct interaction. Functionally, miR-4738-3p counteracted the ability of circ_0074371 to promote M1 macrophage polarization and regulate AMPK phosphorylation. Collectively, these findings indicate that circ_0074371 functions as a miR-4738-3p sponge, providing a novel mechanism underlying its biological activity.

To date, the role of miR-4738-3p in tuberculosis has not been reported. In tumor models, miR-4738-3p has been shown to inhibit tumor cell proliferation and migration^[55]^, whereas in inflammatory diseases, it exerts anti-inflammatory effects^[56]^, These previously reported activities are consistent with the findings of the present study, which also identified an anti-inflammatory role for miR-4738-3p. Moreover, this functional profile supports the proposed mechanism whereby circ_0074371 promotes M1 macrophage polarization by sponging miR-4738-3p.

With respect to the mechanisms underlying macrophage polarization, our findings demonstrate that circ_0074371 promotes M1 macrophage polarization by strongly inhibiting AMPK phosphorylation, thereby exerting a previously unrecognized anti-tuberculosis effect. Previous studies have shown that activation of the AMPK signaling pathway enhances macrophage autophagy and contributes to host defense against Mycobacterium tuberculosis^[15,57,58]^, However, circ_0074371 retained its anti-tuberculosis activity after AMPK inhibition, suggesting that suppression of AMPK by circ_0074371 may activate other antimicrobial immune functions. Evidence from non-tuberculosis models indicates that AMPK activation can attenuate inflammatory responses^[59]^, whereas AMPK inhibition promotes M1 macrophage polarization and the production of inflammatory mediators^[17,60]^, Macrophage polarization is also regulated, at least in part, by the AMPK signaling pathway^[61,62]^. Nevertheless, the role of AMPK in regulating macrophage polarization in tuberculosis models has not been previously described. In the present study, marked inhibition of AMPK activity promoted M1 macrophage polarization and mediated the anti-tuberculosis effects of circ_0074371, revealing a novel AMPK-associated mechanism of tuberculosis resistance. Whether autophagy or macrophage polarization plays the predominant role in host defense against Mycobacterium tuberculosis remains unresolved, and comparative studies addressing this question are lacking. Importantly, these two cellular processes are closely interconnected. For example, M2 macrophage polarization induced by tuberculous pleuritis promotes tumorigenesis through autophagy^[12]^, while autophagy can also alter macrophage polarization phenotypes^[63]^, In the present study, we were unable to determine whether the anti-tuberculosis effects of circ_0074371 in macrophages depend on AMPK signaling. Future studies involving AMPK knockout will be needed to clarify the importance of AMPK in macrophage-mediated anti-tuberculosis immunity and to determine whether AMPK contributes to the remodeling of macrophage polarization.

However, the mechanism by which *MTB* infection downregulates circ_0074371 expression remains unclear. Although this study did not investigate this question in depth, our findings provide an initial direction for future research. Both infection with whole bacteria and stimulation of macrophages with *MTB* components reduced circ_0074371 expression. This suggests that secreted metabolites or intracellular components derived from dead bacteria may act on host genes and thereby affect circ_0074371 expression during infection. Previous studies have shown that *MTB* can modulate host immune responses through secreted molecules^[64]^, Bacterial components may also suppress immunity and promote intracellular survival by impairing host DNA repair^[7]^、altering macrophage metabolism^[65]^and enhancing regulatory T-cell function^[66]^, In mouse models, *MTB* produces D-serine to limit CD8+ T-cell-dependent immune responses^[67]^; The host, in turn, can suppress immune responses by modifying protein components derived from *MTB* ^[68]^, Taken together, these findings raise the possibility that secreted *MTB* proteins affect circ_0074371 circularization, thereby reducing its production. Future studies will identify the bacterial components that target this circularization mechanism, with the aim of developing an additional tuberculosis treatment strategy directed at these components.

Because of their greater stability and biocompatibility than linear RNAs, circRNAs may be particularly suitable for development as bioactive therapeutic agents. Preliminary studies have explored the use of similar molecules as immunomodulators for tuberculosis treatment^[30]^. Our findings expand the regulatory network through which circRNAs modulate host immunity and identify circ_0074371 as a potential immunotherapeutic target for tuberculosis control.

## Acknowledgments

This work was supported by the National Natural Science Foundation of China (81870016, 82270013), the Guangdong Basic and Applied Basic Research Foundation (2022A1515140190, 2021B1515140068), the Science and Technology Project of Dongguan (20211800904782, 20221800905862), Science and Technology Program of Shenzhen (JCYJ20220530144402005),“Zhumeng Yangfan” Open Collaborative Research Foundation of Shenzhen Bao’an District Songgang People’s Hospital (External Cooperation Project: SGYYZ2026B-08), “Qilin Talent” Post Scientific Research Start-up Project of Shenzhen Bao’an District Songgang People’s Hospital(SGYYQ2026A001), Shenzhen Bao’an District Medical and Health Scientific Research Projec(t 2026JD089), the Special Youth Cultivation Project of the Dongguan Basic Research Fund (20251800400912) and the National College Students’ Innovation and Entrepreneurship Training Program(202410571005).

---

**Figure S1.**
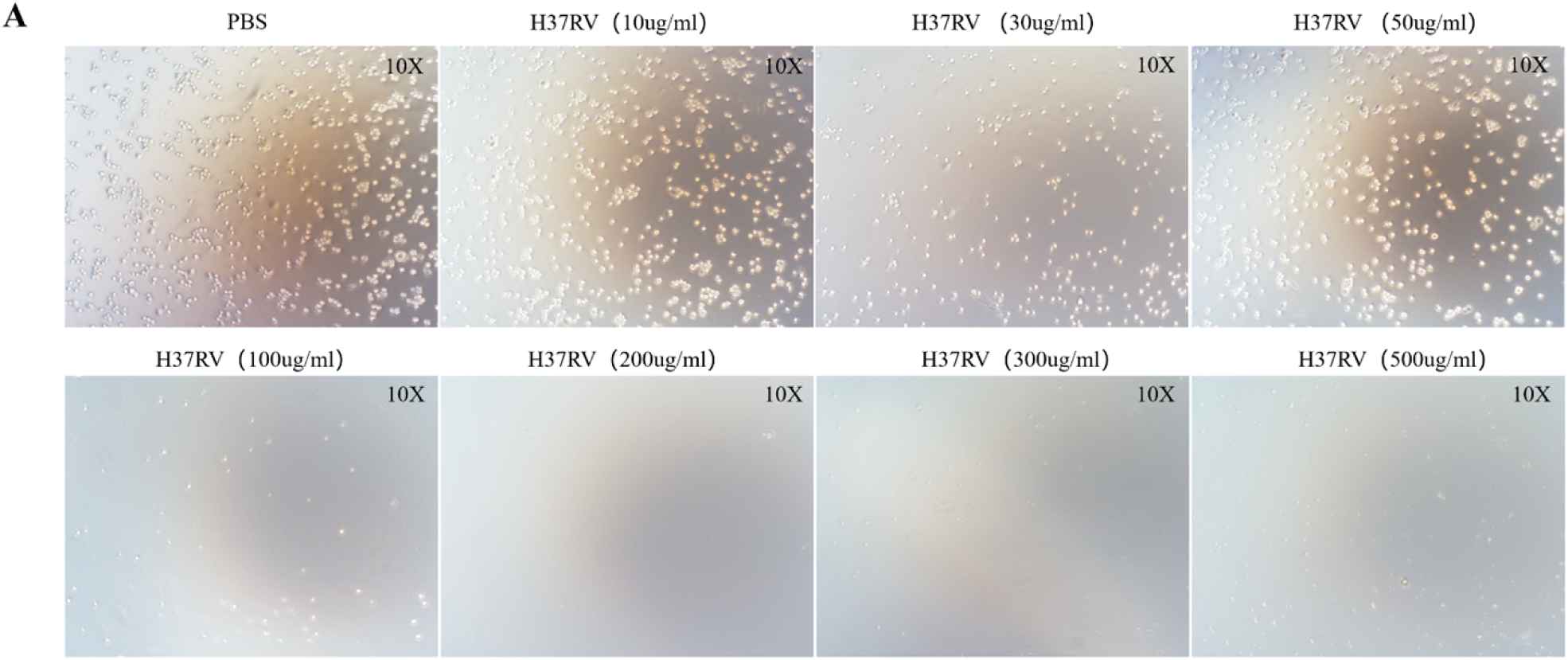
Microscopic morphology of THP-1 cells following treatment with varying concentrations of *H37Rv* lysate

**Figure S2.**
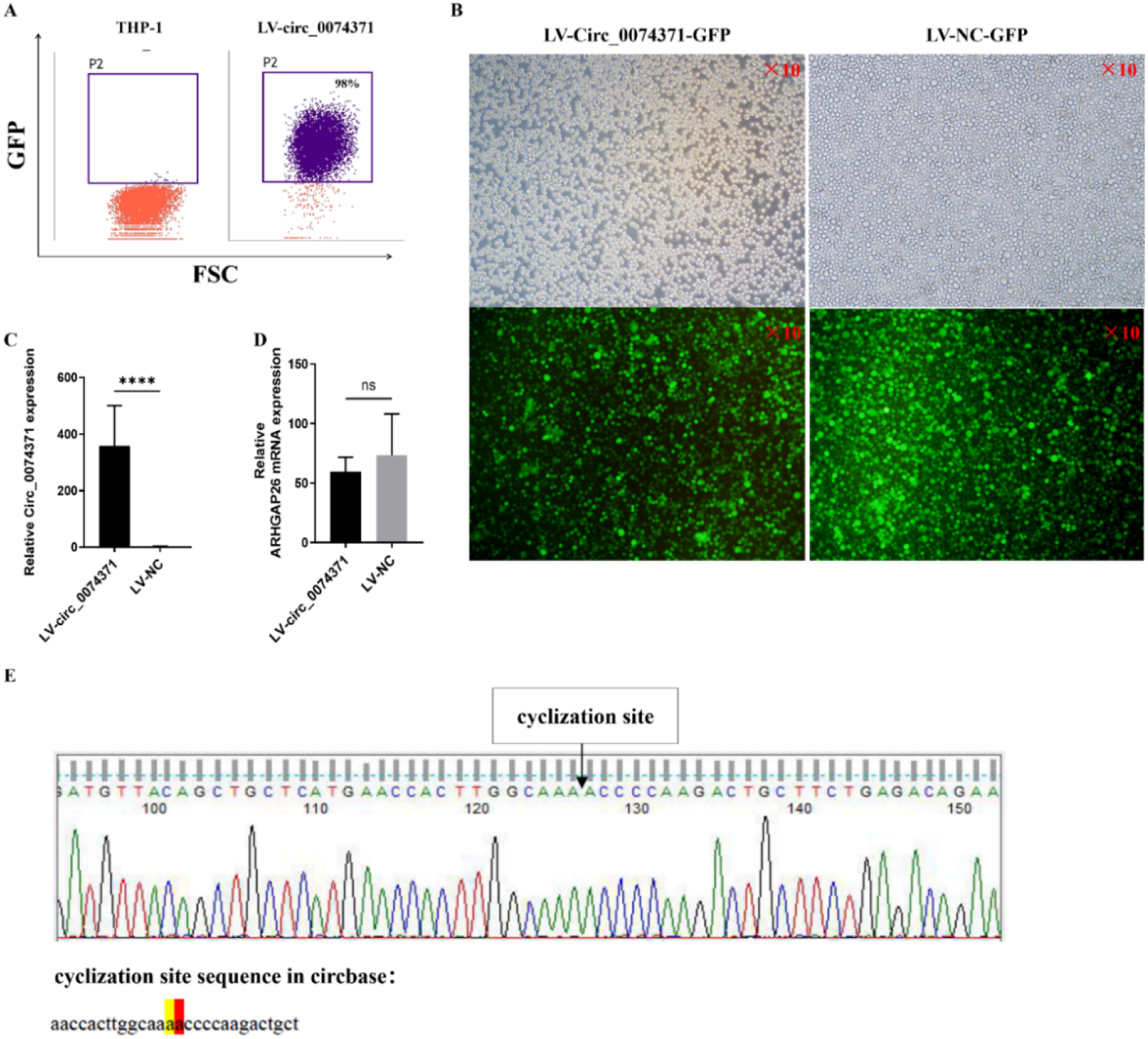
Generation of circ_0074371-overexpressing cell lines using lentiviral vectors (A) Flow cytometric analysis of lentiviral transduction efficiency. (B) Fluorescence microscopy assessment of lentiviral transduction efficiency. (C) RT-PCR validation of circ_0074371 overexpression. (D) RT-PCR analysis of the effect of circ_0074371 overexpression on host gene ARHGAP26 expression. (E) Sanger sequencing confirmation of the back-splice junction sequence of circ_0074371.

**Figure S3.**
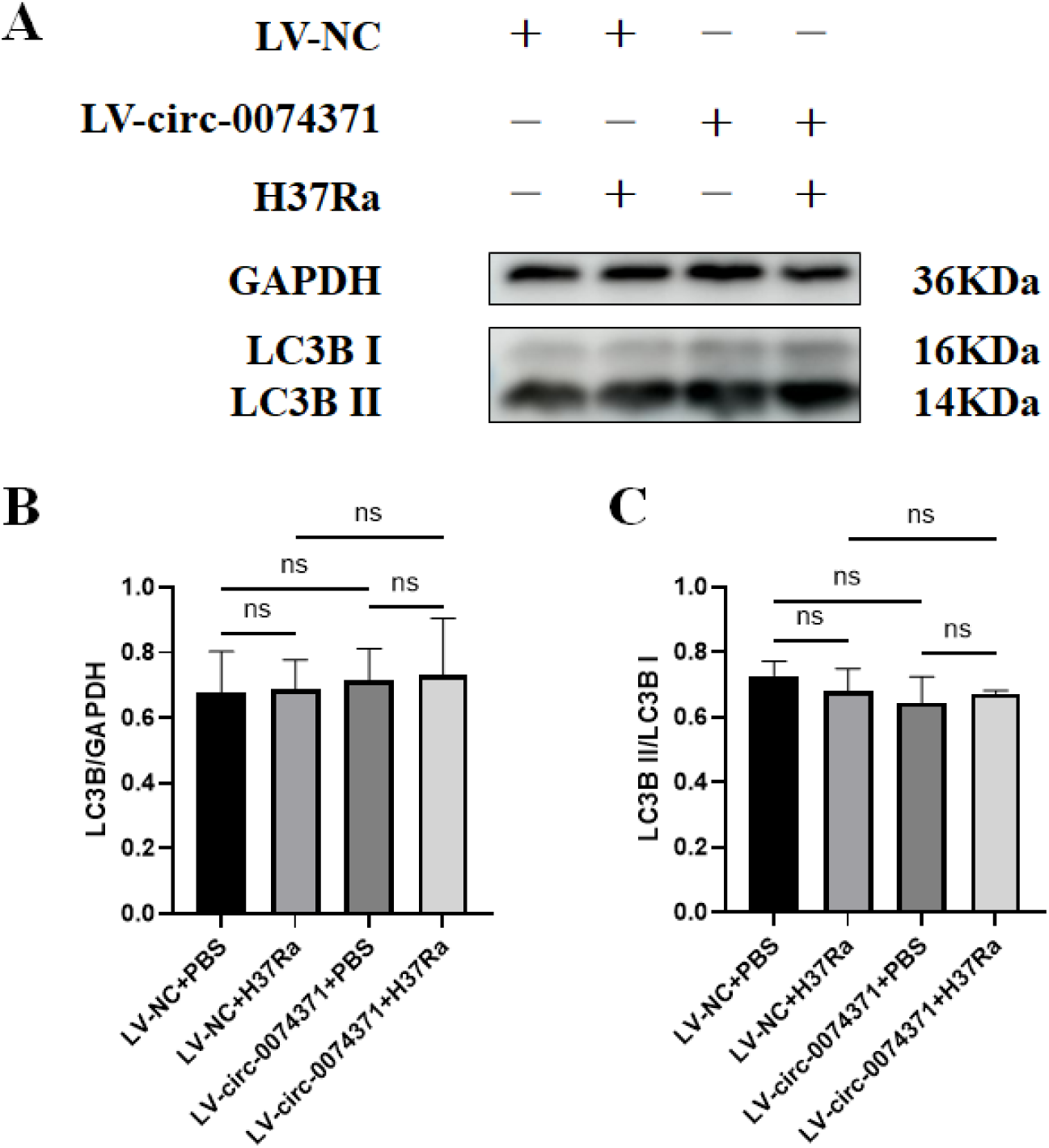
Circ_0074371 has no effect on macrophage autophagy. (A) THP-1 cells overexpressing circ_0074371 were infected with *Mycobacterium tuberculosis H37Ra* for 24 h, after which autophagy markers LC3B-I and LC3B-II were assessed by Western blotting. (B) Quantification of LC3B expression normalized to GAPDH. (C) Quantification of the LC3B-II/LC3B-I expression ratio.

**Table S1.** Clinical Sample Information Statistics for circRNA Detection.

|  | ATB | HC | *P* |
| --- | --- | --- | --- |
| Age(Y) | 35.90±9.54 | 36.47±10.63 | >0.05 |
| Gender (F/M) | 16/15 | 19/ 17 | >0.05 |
| T(M0/M1) | 31/16 | 36/0 | - |

**Table S2.** Clinical Sample Information Statistics for miRNA Detection.

|  | ATB | HC | *P* |
| --- | --- | --- | --- |
| Age(Y) | 37.31±10.86 | 38.4±8.67 | >0.05 |
| Gender (F/M) | 5/11 | 6/ 9 | >0.05 |
| T(M0/M1) | 16/0 | 15/0 | - |

**Table S3.** Clinical characteristics of the enrolled subjects.

| ATB |  |  |  |  |  | HC |  |  |  |
| --- | --- | --- | --- | --- | --- | --- | --- | --- | --- |
| No | Age | Sex(F/M) | T (d) | IT/RT | Circ/miRNA | No | Age | sex (F/M) | Circ/miRNA |
| 1 | 51 | M | 0, 30 | IT | Circ | 1 | 41 | F | Circ |
| 2 | 25 | F | 0, 30 | IT | Circ | 2 | 34 | M | Circ |
| 3 | 38 | M | 0, 30 | IT | Circ | 3 | 52 | F | Circ |
| 4 | 52 | M | 0, 30 | IT | Circ | 4 | 27 | M | Circ |
| 5 | 38 | F | 0, 30 | IT | Circ | 5 | 34 | F | Circ |
| 6 | 26 | F | 0, 30 | IT | Circ | 6 | 33 | M | Circ |
| 7 | 47 | F | 0, 30 | IT | Circ | 7 | 47 | M | Circ |
| 8 | 38 | M | 0, 30 | IT | Circ | 8 | 22 | M | Circ |
| 9 | 30 | F | 0, 30 | IT | Circ | 9 | 45 | F | Circ |
| 10 | 37 | F | 0, 30 | IT | Circ | 10 | 41 | F | Circ |
| 11 | 41 | M | 0, 30 | IT | Circ | 11 | 32 | F | Circ |
| 12 | 25 | F | 0, 30 | IT | Circ | 12 | 38 | M | Circ |
| 13 | 41 | F | 0, 30 | IT | Circ | 13 | 42 | M | Circ |
| 14 | 28 | M | 0, 30 | IT | Circ | 14 | 28 | F | Circ |
| 15 | 33 | M | 0, 30 | IT | Circ | 15 | 39 | M | Circ |
| 16 | 53 | M | 0, 30 | IT | Circ | 16 | 41 | M | Circ |
| 17 | 32 | F | 0 | IT | Circ | 17 | 36 | M | Circ |
| 18 | 27 | F | 0 | IT | Circ | 18 | 45 | F | Circ |
| 19 | 33 | F | 0 | IT | Circ | 19 | 48 | F | Circ |
| 20 | 23 | M | 0 | IT | Circ | 20 | 55 | F | Circ |
| 21 | 32 | F | 0 | IT | Circ | 21 | 24 | M | Circ |
| 22 | 29 | M | 0 | IT | Circ | 22 | 45 | F | Circ |
| 23 | 32 | F | 0 | IT | Circ | 23 | 45 | M | Circ |
| 24 | 50 | M | 0 | IT | Circ | 24 | 56 | M | Circ |
| 25 | 29 | M | 0 | IT | Circ | 25 | 33 | M | Circ |
| 26 | 52 | F | 0 | IT | Circ | 26 | 19 | F | Circ |
| 27 | 47 | F | 0 | IT | Circ | 27 | 25 | F | Circ |
| 28 | 29 | M | 0 | IT | Circ | 28 | 34 | F | Circ |
| No | Age | Sex(F/M) | T (d) | IT/RT | Circ/miRNA | No | Age | sex (F/M) | Circ/miRNA |
| 29 | 20 | F |  | IT | Circ | 29 | 28 | M | Circ |
| 30 | 42 | M |  | IT | Circ | 30 | 55 | F | Circ |
| 31 | 33 | M |  | IT | Circ | 31 | 24 | F | Circ |
| 32 | 34 | F |  | IT | miRNA | 32 | 33 | M | Circ |
| 33 | 35 | F |  | IT | miRNA | 33 | 18 | F | Circ |
| 34 | 43 | M |  | IT | miRNA | 34 | 46 | F | Circ |
| 35 | 22 | M |  | IT | miRNA | 35 | 25 | M | Circ |
| 36 | 28 | M |  | IT | miRNA | 36 | 23 | F | Circ |
| 37 | 45 | M |  | IT | miRNA | 37 | 41 | M | miRNA |
| 38 | 38 | M |  | IT | miRNA | 38 | 44 | M | miRNA |
| 39 | 33 | M |  | IT | miRNA | 39 | 45 | M | miRNA |
| 40 | 28 | M |  | IT | miRNA | 40 | 46 | F | miRNA |
| 41 | 39 | F |  | IT | miRNA | 41 | 33 | F | miRNA |
| 42 | 27 | M | 0 | IT | miRNA | 42 | 50 | M | miRNA |
| 43 | 53 | F | 0 | IT | miRNA | 43 | 39 | M | miRNA |
| 44 | 31 | F | 0 | IT | miRNA | 44 | 26 | F | miRNA |
| 45 | 50 | M | 0 | IT | miRNA | 45 | 47 | M | miRNA |
| 46 | 29 | M | 0 | IT | miRNA | 46 | 29 | F | miRNA |
| 47 | 62 | M | 0 | IT | miRNA | 47 | 35 | M | miRNA |
|  |  |  |  |  |  | 48 | 30 | F | miRNA |
|  |  |  |  |  |  | 49 | 35 | M | miRNA |
|  |  |  |  |  |  | 50 | 51 | F | miRNA |
|  |  |  |  |  |  | 51 | 25 | M | miRNA |

**Table S4.**
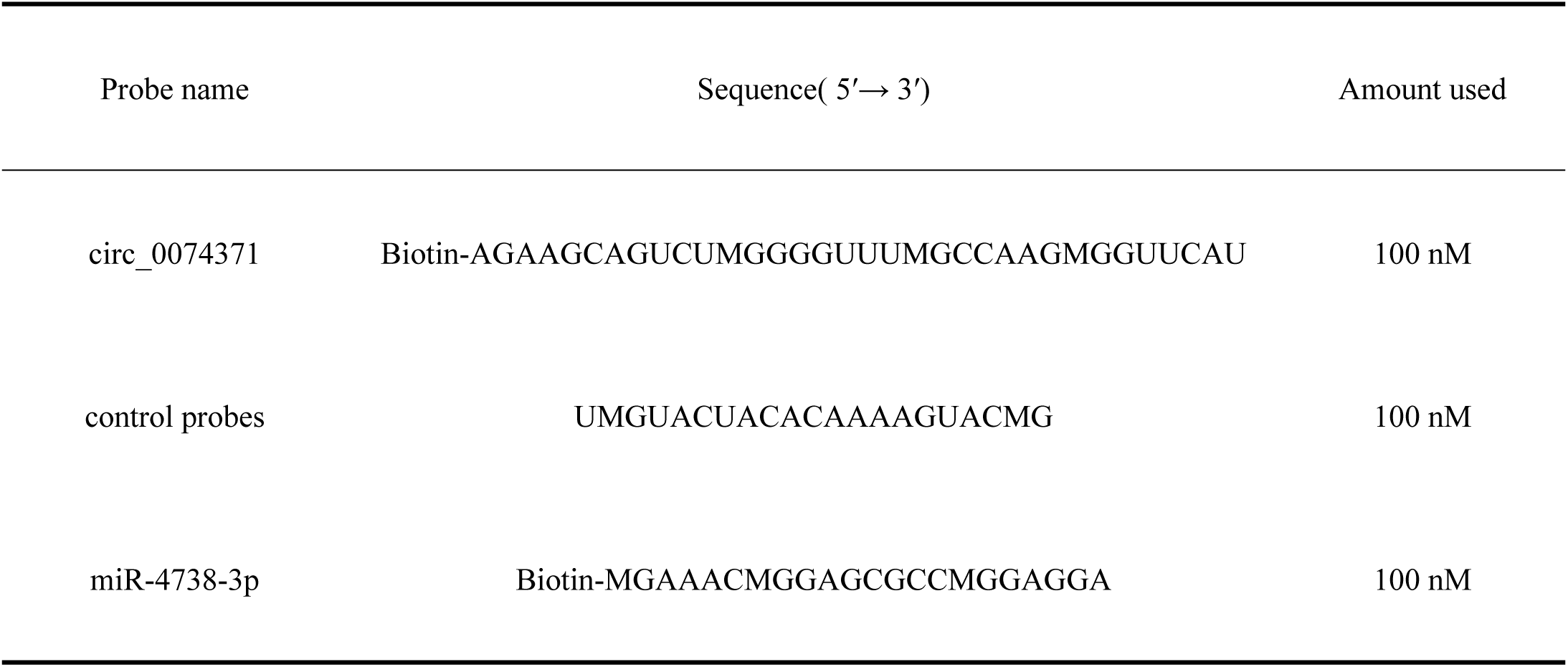

**Table S5.**
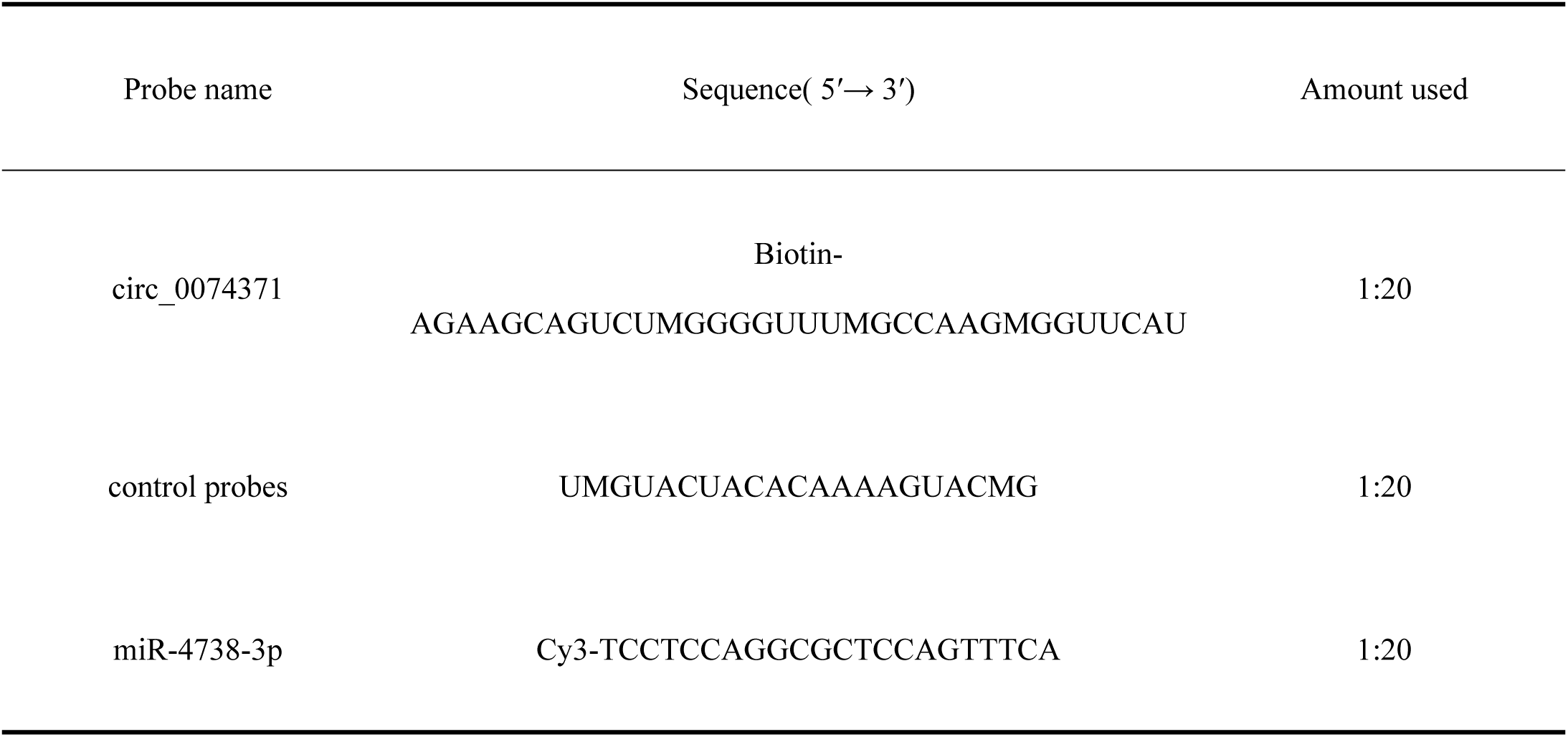

